# Eye-specific detection and a multi-eye integration model of biological motion perception

**DOI:** 10.1101/2023.09.24.559175

**Authors:** Massimo De Agrò, Daniela C. Rößler, Paul S. Shamble

## Abstract

The term biological motion refers to the peculiar kinematics of living organisms. Their interconnected joints move at a fixed distance from each other, a pattern that is common among all locomotive, rigid animals. Across the animal kingdom, many species have developed specialized circuitry to visually recognize biologically moving stimuli and discriminate them from other patterns. Recently, this skill has also been observed in the distributed visual system of jumping spiders. These eight-eyed animals use three of their eye pairs to perceive motion. Then, the gaze of the remaining pair is shifted towards the detected object for further inspection. When presented with a biologically moving stimulus and a random one, jumping spiders turn to face the latter, demonstrating discrimination. In the current paper, we systematically tested the ability of jumping spiders to discriminate biological from random displays using every single eye-pair, by blocking the others with paint. The animals were able to discriminate the stimuli only when the anterior-lateral eyes were unblocked, performing at chance level with the other pairs. Crucially, the spiders preferred the biological stimulus, not the random one. To explain this preference reversal we hypothesized a model, describing how the anterior-lateral eyes’ specialization in detecting biological motion feeds into a multi-eye integration system, generating more complex behavior from the combination of the simple, single-eye responses. We propose that this in-built modularity may be a solution to the limited resources of these invertebrates’ brains, constituting a novel approach to visual processing.

## 1 Introduction

Many animals have photosensitive cells, that enable them to acquire visual information from the environment (Lazareva et al., 2012). Collecting light, however, is not always enough—for more sophisticated visual tasks, patterns of activation need to be organized and interpreted to inform subsequent decision making (DiCarlo et al., 2012). Due to the high variety of visual information, this is a complex task. In humans and other vertebrates, this has driven the evolution of massive neural networks that use hierarchical processes to interpret the visual scene (Felleman & Van Essen, 1991; Hubel & Wiesel, 1962; Serre, 2014; Van Essen et al., 1992). Indeed, increasing neuronal investment by enlarging the brain, as it is the case in humans, seems to be an effective solution to the problem of complexity (Hofman, 2014). This however is not an option for smaller animals, as they need to accomplish many of the same visual tasks, but don’t have the space to grow their brain. Arthropods seem to have found a solution. Behavioral evidence suggests that the comparatively small size of their nervous system does not hinder the production of complex behaviors—including many that are comparable to those of vertebrates (Avarguès-Weber et al., 2012; Avarguès-Weber & Giurfa, 2013; Chittka & Niven, 2009; De Agrò et al., 2020; Eberhard, 2007, 2011; Eberhard & Wcislo, 2011; Goté et al., 2019; Wehner, 2003).

Jumping spiders, in particular, show an impressive level of cognitive and behavioral complexity despite having a relatively small nervous system. These include learning (De Agrò, 2020; De Agrò et al., 2017; Liedtke & Schneider, 2014; Mannino et al., 2023), numerical abilities (Cross & Jackson, 2017), spatial and action planning (Cross & Jackson, 2015, 2016, 2019; Tarsitano & Jackson, 1994), object recognition (Dolev & Nelson, 2014, 2016; Rößler, De Agrò, et al., 2022), and even engaging in REM sleep-like behaviors (Rößler, Kim, et al., 2022). Fittingly, the most impressive characteristic of these animals is their vision (Winsor et al., 2023): a modular, specialized system split into 4 pairs of eyes (Figure 1), with each pair projecting into anatomically separate brain areas (Harland et al., 2012; N. Morehouse, 2020; N. I. Morehouse et al., 2017; Steinhoff et al., 2017, 2020). The two largest, forward-facing, anterior medial eyes (AMEs, principal eyes, Figure 1) have the highest visual acuity and a layered retina that allows for single-eye depth perception and color vision (Land, 1969b; Nagata et al., 2012; Zurek et al., 2015). These eyes possess a narrow visual field (≈5°), that can be shifted around thanks to a set of muscles, (Land, 1969a)—achieving a similar function as the fovea in human eyes. The other three pairs of eyes—the anterior lateral eyes (ALEs), posterior median eyes (PMEs) and posterior lateral eyes (PLEs) are collectively called secondary eyes (Figure 1). They are monochrome and smaller, but possess a much wider visual field with a combined range of ≈350°. The principal and secondary eyes divide the labor of visual computation (Strausfeld et al., 1993; Strausfeld & Barth, 1993), with the primary eyes handling static detail while the secondary eyes specialized in perceiving movement. As soon as a stimulus is detected, the spider rapidly pivots in order to face it with the principal eyes (Beydizada et al., 2023; Ferrante et al., 2023; Zurek et al., 2010; Zurek & Nelson, 2012a). The AMEs can then start scanning the object, with the task of recognizing it (Dolev & Nelson, 2014, 2016; Land, 1969a; Menda et al., 2014; Rößler, De Agrò, et al., 2022; Zurek & Nelson, 2012b).

**Figure 1.**
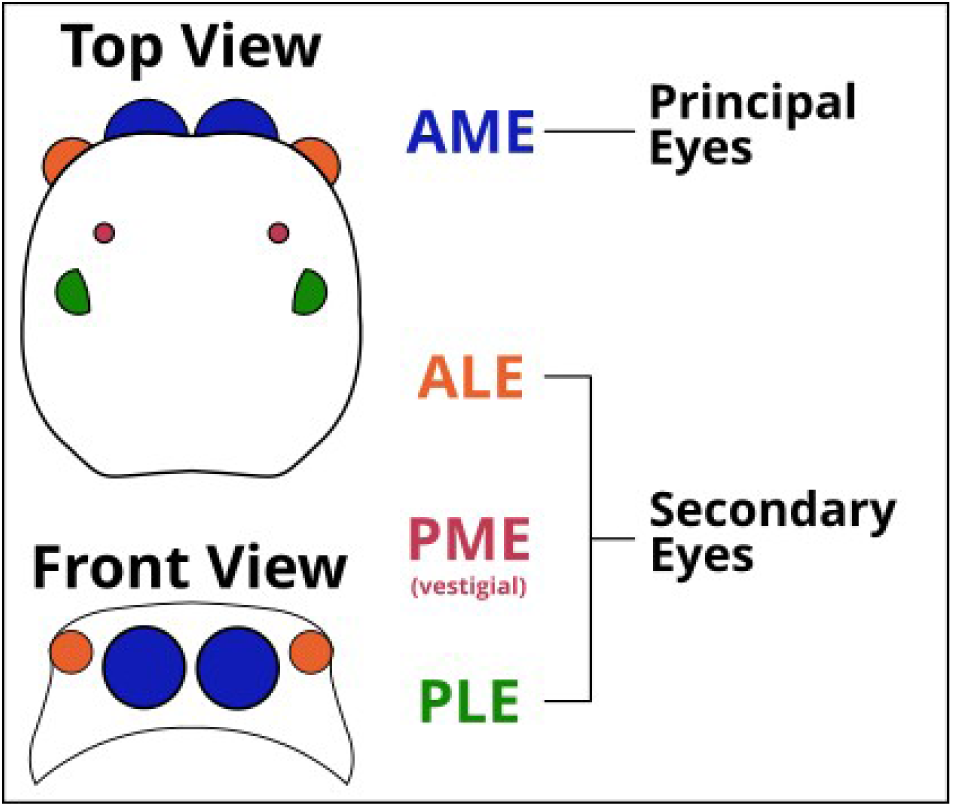
Schematic representation of the 8 eye pairs of the jumping spider, top and front view. The Anterior Medial Eyes (AMEs) are named Principal Eyes and are movable, with a small visual field of around 5° each, but with high spatial resolution and color vision. These eyes are likely specialized for static figure discrimination. The other three pairs of eyes, Anterior Lateral (ALE), Posterior Medial (PME) and Posterior Lateral (PLE) are named Secondary Eyes. With a wider visual field but lower acuity, they are considered to be specialized for motion perception and discrimination.

The jumping spiders’ secondary eyes are not limited to detecting moving objects, but can also recognize different types of motion and can inform subsequent behaviors accordingly (Bruce et al., 2021; De Agrò et al., 2021; Spano et al., 2012). This ability also extends to quite complex dynamic visual stimuli. In a previous experiment (De Agrò et al., 2021) we demonstrated that jumping spiders can recognize biological motion—a term that refers to stimuli that move according to a pattern common to all living organisms (Johnson, 2006). During experiments, these stimuli are generally presented as clouds of dots, with motion that maps an animals’ major joints during locomotion, yet these cloud patterns are devoid of any geometrical structure (Johansson, 1973, 1976; Lemaire & Vallortigara, 2022; Neri et al., 1998; Troje, 2013; Troje & Westhoff, 2006).

To date, it is not clear whether the AMEs are required to perform this recognition. Moreover, it is still unknown whether all secondary eyes can perform this discrimination or if it is specialized to a single eye-pair. We hypothesize that due to the selection for functional specialization in the visual systems of jumping spiders, biological motion detection occurs in a single eye-pair—most likely the ALEs, as these eyes are motion-sensitive and forward-facing—rather than being distributed across multiple eyes and brain areas. Note that to discriminate biologically moving stimuli from random moving ones, it is not sufficient to analyze a single dot trajectory—rather, one must integrate the relative motions of multiple entities. In humans and other vertebrates this complex integration is carried out by late visual areas, like the medial temporal area MT (E. Grossman et al., 2000; E. D. Grossman & Blake, 2002). The presence of discrimination in a single eye-pair would suggest that the computation happens in a dedicated brain area, very early in the visual stream, suggesting the presence of a currently unknown process to detect biological motion.

To test this hypothesis, we selectively covered jumping spiders’ eyes to leave them either with only ALEs, only PLEs, or both pairs of secondary eyes un-occluded. We then presented animals with pairs of stimuli, one of biological motion and the other depicting random motion, and recorded which of these stimuli they turned towards.

## 2 Materials and methods

### 2.1 Subjects

For the experiments, we collected jumping spiders of the species *Menemerus semilibatus* from the field. These spiders can be found in parks and on buildings and are abundant across southern Europe. For the first experiment, we used 31 individuals (9 males, 14 females, 8 juveniles) 179 were used for the second, main experiment (15 males, 89 females, 75 juveniles). Only animals with a body length over 5 mm were collected to ensure functioning of the methodology. Once caught, the animals were housed in clear plastic boxes (dimensions 80 × 65 × 155 mm). They were fed *Drosophila* fruit flies *ad libitum*, replenished once a week, until used for the experiment. The day before testing, a magnet was fixed to the prosoma (head) of each subject using UV glue (brand) in order to attach the animal to the treadmill apparatus (see De Agrò et al., 2021).

Concurrently, we covered the spider’s eyes according to their assigned experimental treatment. Each spider was assigned to one of following three treatments:

- ALE treatment – The animal had only their ALEs uncovered, as paint was applied over AMEs, PMEs and PLEs.
- PLE treatment – The animal had only their PLEs uncovered, with paint applied over AMEs, ALEs and PMEs.
- ALE+PLE treatment – The animal had ALEs, PMEs and PLEs uncovered, with paint applied only over AMEs.

We did not include a PME condition. PMEs are believed to be vestigial with a limited field of view (Land, 1985; see Figure 1 for the eyes organization).

Spider eyes were painted under the microscope using a toothpick with a small dab of water-based white paint. White paint was chosen over other colors as it was easier to see the coverage on the surface of the dark-colored spider eyes. After each animal underwent all the trials it was assigned to, the magnet was removed, and the paint washed off. The spider was then released in the same spot where it was captured. Magnets did not appear to negatively affect the animals during the short period in which they were housed in the lab, and spiders freed off the magnet appeared to move and behave normally.

### 2.2 Experimental apparatus

The experimental apparatus, stimuli, and scoring used in these experiment were the same as the ones described in De Agrò et al. (2021), except that the computer monitors were arranged differently (see below). In brief, a polystyrene sphere (38mm diameter) was caged in a plastic holder, suspended by constant stream of compressed air, and positioned in the center of the apparatus. The top of the plastic holder was open, revealing a 20mm wide cap of the sphere below. At the start of each experiment a spider was attached to the end effector of a 6 axis micro-manipulator using the magnet glued on top of the prosoma. Then, the animal was lowered and positioned with the legs in contact with the polystyrene sphere and oriented relative to the monitors. This way, despite being tethered, the spider was able to affect motion of the sphere below. By recording the sphere using a high-speed camera (120fps), we extracted the frame-by-frame rotational matrices (Moore et al., 2014), and thus, the intended motion of the spider. Two computer monitors (BenqGW2270, 1920x1080, 537mm wide) were placed in front of the animal, angled to each other at 120*°* or 65*°*, depending on the experimental condition (see below). The contact point between the two monitors was placed directly in front of the animal, at the center of their visual field, from here on defined as 0°. In both experiments stimuli appeared in one or both of the two monitors, moving from the outermost section towards the center, or vice-versa. When detecting a stimulus with the secondary eyes, jumping spiders performed full-body pivots (Land, 1972), to focus the AMEs visual fields on the target. When presented with opposing information—for example, stimuli on two different sides—the spider preferentially turned towards one depending on valence, preference, or possibly other factors (De Agrò et al., 2021). In the current experiment, we recorded the pivots performed by animals towards stimuli presented on the monitors (see Figure 2), to infer either detection angle (when only one was presented, see Experiment 1) or relative inspection preference (when two were presented simultaneously, see Experiment 2).

**Figure 2.**
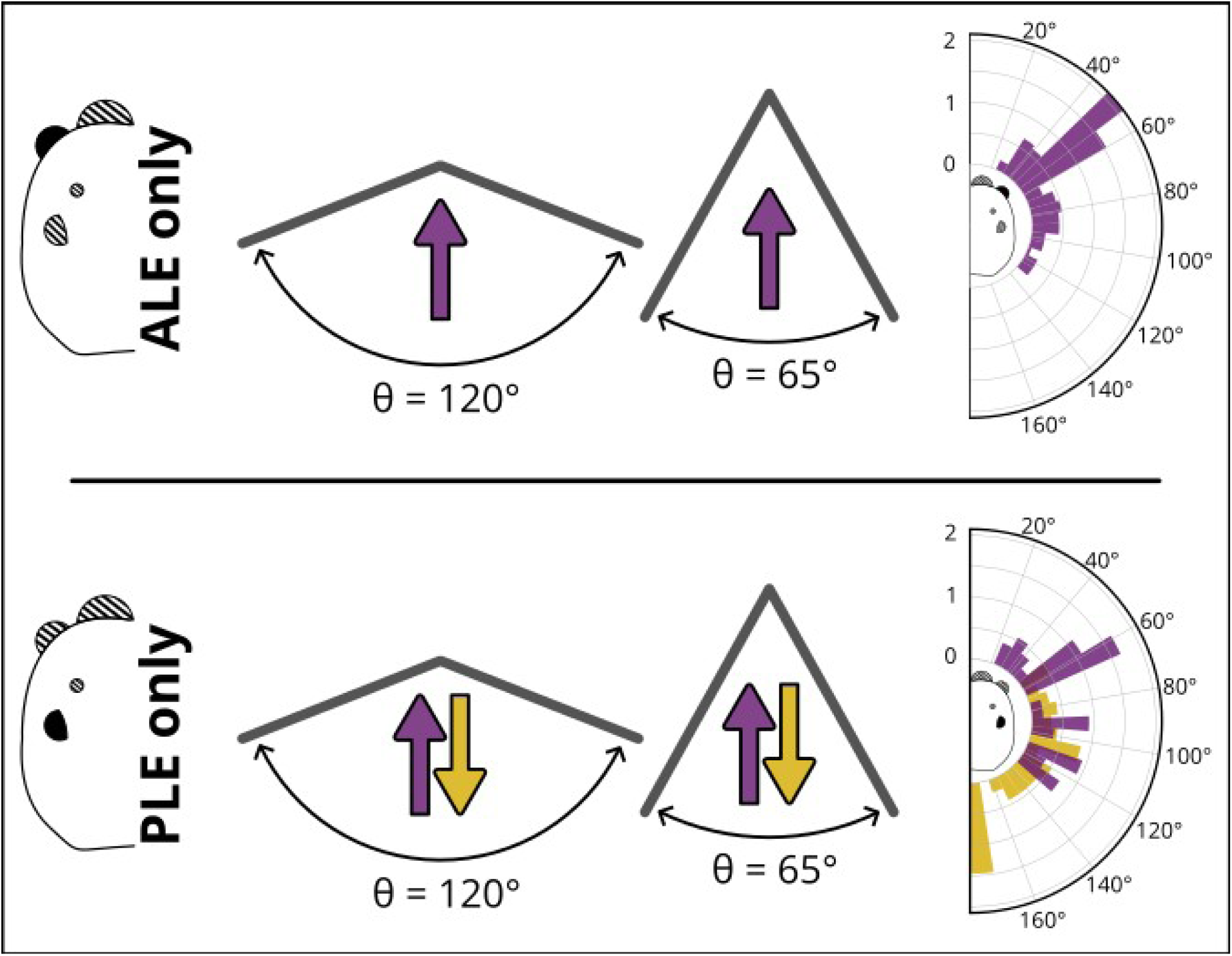
Procedure and results for both conditions of experiment 1. In the ALE only condition, the spiders were placed on two different setups: facing the meeting point of two monitors, angled relative to each other either 120° or 65° . Facing direction indicated by the purple arrow. The graph on the right reports the relative frequency of saccade per position of the moving stimulus. Y-axis=relative frequency of saccade. X-axis=angle of the stimulus. A clear peak in rotation frequency is appreciable at 50°. In the PLE only condition, the spiders were placed in the same two setups, but they could be oriented either towards the meeting point (purple) or away from it (gold). This was done to be able to present stimuli all around the animal, rather than only in the front. In the result graph, purple bars are for trials where the spiders were oriented towards the screens, gold bars are for when the spiders were backward (the graph is still represented from the spider’s point of view, hence the bars at the spider’s back). When oriented forward, a clear peak is appreciable at 60°. When oriented backwards, the peak is visible at 180°.

### 2.3 Experiment 1 – Identification of eye-specific visual angles

In the first experiment, we used a behavioral procedure to identify the extent of the visual field of every eye-pair in *Menemerus semilimbatus.* Information about eye-specific visual field spans is available in the literature for other jumping spider species (Land, 1985; Zurek & Nelson, 2012b), but visual fields vary considerably across different species of Salticidae (Land, 1985). To draw this visual field map, we exploited the spiders’ typical secondary eyes detection behavior described in the introduction: since these animals tend to perform a pivot immediately when a moving stimulus is detected, an object moving horizontally will trigger a reaction from the spider as soon as it enters its visual field. By recording the angular position of the stimuli upon first detection, we can behaviorally draw the edges of every eye.

In order to present stimuli across the entire 360° around the animal, we varied the placement of the monitors and of the spiders across the experiment (Figure 2). First, the two monitors were placed angled to each other either at 120° or 65°. The spider was positioned at a distance of 200 mm from the contact point of the two monitors. The different position of the two monitors of course allowed for a different coverage of the spiders’ visual field, with the first option spanning across ≈200° and the second ≈265°. To cover the remaining ≈95° at the back of the spider, we decided to reverse the spiders’ orientation, facing opposite to the monitors meeting points (Figure 2).

Each spider in the PLE treatment (n=16) underwent 4 conditions across 4 randomly ordered trials, one for each setup (monitors at 120°, frontally facing; monitors at 65°, frontally facing; monitors at 120°, backward facing; monitors at 65°, backward facing. Figure 2). Spiders in the ALE treatment (n=15) instead underwent 2 conditions, corresponding to the two frontally facing setups (Figure 2), as the ALEs visual field is generally identified as being ±50° (Zurek & Nelson, 2012b). No ALE+PLE spider underwent experiment 1, as the combined visual field would have not been informative.

At the start of each trial, each spider was positioned on the polystyrene sphere, oriented according to the given trial and condition. After 210 seconds of habituation, a stimulus appeared either on the left or the right monitor, starting either from the contact point of the two monitors or the outer border. The stimulus then moved at 9°/s (the speed previously shown to trigger the highest saccadic probability. See Zurek et al., 2010), either towards the center of the visual field or away from it, until it reached the opposing edge of the monitor. A new stimulus appeared 15 seconds later, for a total of 30 presentations. For each presentation, we recorded the first pivot produced by the spider, and noted the angle position of the stimulus at pivot initiation.

### 2.4 Experiment 2 – Eye-specific preference for biological motion

In experiment 1, trials in which the monitors were positioned at 65° elicited the highest number of responses. Moreover, we observed the greatest number of responses in the ALE treatment when the stimuli were located at ±50°. For the PLE treatment, we observed peaks in responses at ±60° and ±180° (for the full description, see Results and SM02. Figure 2). Consequently, we positioned monitors at 65° for Experiment 2, with the spider oriented towards the monitors, thereby covering both the ALEs and PLEs field of views, and most importantly the crossing point between the two.

Each subject (n=179) was assigned either to the ALE+PLE (N=61), ALE (N=58), or PLE (N=60) treatment. The spider then underwent two different conditions, across two trials performed in the same day. In the first condition, the spider was positioned on the spherical treadmill and allowed to habituate for 210 seconds. Then, two stimuli—a random point-light display and a biological point-light display—appeared at ±90° and moved towards the center of the screen. These stimuli are the same used in our previous experiment (De Agrò et al., 2021). The two stimuli proceeded with the same speed and occupied the same angular position to each other in each frame. The stimuli then stopped at ±50° for one second before starting to move again until they disappeared near the contact point of the two monitors (point of disappearance = ±10°). The second condition proceeded identically to the first, but the stimuli presented were a moving spider silhouette and an equally sized ellipse (taken from De Agrò et al., 2021 as well). The order of these two conditions was randomized for each spider.

After each stimulus pair, there was a pause of 25 seconds before a second pair appeared, for a total of 20 repetitions per trial. For each repetition, the position (left/right) of the two stimuli was randomized, as well as the movement direction (either both moving inward, from ±90° towards ±10°, or outwards, from ±10° towards ±90°).

### 2.5 Scoring

The frame-by-frame rotation of the sphere was extracted using the software FicTrac (Moore et al., 2014). We then focused on the sphere rotations around its Z-axis, as they correspond to the spiders’ pivots. Using a custom script written in Python 3 (Van Rossum & Drake, 2009), and the packages *pandas* (Jeff Reback et al., 2020), *numpy* (Oliphant, 2006; van der Walt et al., 2011) and *scipy* (Virtanen et al., 2020), we detected peaks in the signal, corresponding to probable rotation events. Positive and negative peaks were recorded, as they correspond to clockwise and counterclockwise rotations. Each peak could then be associated with the position of the stimulus based on the time of stimulus appearance and the recorded time of the peak. The full script is available as a supplement.

For Experiment 1, we were interested in the first pivot performed by the spider, which most likely indicates when the stimuli first enter the animal’s visual field. Therefore, we selected the first measured peak with a rotation minimum of 20° (determined by calculating the area under the curve for the selected signal peak) in the direction of the stimulus (clockwise for stimuli on the left, counterclockwise for stimuli on the right—note that the spider’s intended rotation is opposite to the rotation of the sphere). We then recorded the position of the stimulus at the time of the first rotation and saved it as the point of detection.

As Experiment 2 followed almost the same procedure as our previous work (De Agrò et al., 2021), we followed the same procedure for the scoring. In brief, after selecting all the peaks in the Z-axis as before, we changed their sign according to the biological stimulus position. This way, rotations in the direction of the biological (or silhouette) stimulus were set as positive values, while rotation in the direction of the random (or ellipse) stimulus were set as negative. All peaks were then included in the analysis, to compute a general pivot tendency. If the spiders performed an equal number of pivots towards the biological and the random stimulus, this would result in an average approaching 0. Likewise, an average > 0 would corresponded with a preference for the biological stimuli/silhouette, while an average < 0 would corresponded with a preference for the random/ellipse. To confirm the validity of this scoring procedure, we also applied it to the results of Experiment 1. Here, only one stimulus at a time was available, and as such having no other target the spiders were all expected to turn towards it, resulting in an average significantly and consistently higher than 0.

### 2.6 Statistical Analysis

All analyses were performed using R 4.2.1 (R Core Team, 2020), including the libraries *readODS* (Schutten et al., 2020), *glmmTMB* (Brooks et al., 2017; Magnusson et al., 2020), car (Fox & Weisberg, 2019), *DHARMa* (Hartig, 2020), *emmeans* (Lenth, 2021), *ggplot2* (Wickham et al., 2020) and *reticulate* (Ushey et al., 2021). Graphical outputs were produced using Python 3 (Van Rossum & Drake, 2009, p. 3), with the packages *matplotlib* (Hunter, 2007) and *seaborn* (Waskom et al., 2017).

We employed generalized linear models in our analysis. For each model we included the subject identity as a random intercept and experimental condition as random slope—as different subjects could have both a different base reactivity (intercept) and a differential response to the conditions (slope). However, this resulted in over-fitting in some cases, which prompted us to remove condition as a random slope. For Experiment 1, we modeled the saccadic probability as influenced by treatment (ALE, PLE) and monitor setup (120°, 65°) using a binomial error structure. Regarding the angle of first detection, we plotted the relative frequencies of rotation against the angle of the stimulus and derived the section of highest reactivity. For Experiment 2, we modeled the z-rotation speed as influenced by treatment (ALE, PLE, ALE+PLE) and condition using a Gaussian error structure.

Below we will report only the main findings. For the full analysis and raw data please refer to SI.

## 3 Results

### 3.1 Experiment 1 – Identification of eye-specific visual angles

As anticipated, we observed a higher response probability for the 65° screen orientation versus the 120° orientation (GLMM post-hoc, Bonferroni corrected; odds.ratio=2.75, SE=0.547, t=5.084, p<0.0001), with no significant difference between ALE and PLE treatments (odds.ratio=2.28, SE=1.154, t=1.629, p=0.207).

When considering the position of the stimulus upon pivot initiation, there was a clear peak in responses at around ±50° for the ALE treatment (Figure 2). This is consistent with published values for other species (Zurek & Nelson, 2012b), and suggests the total visual span of the ALEs to be ≈100°. For PLE treatment, when the animals were oriented frontally the majority of responses occurred at around ±60°. When the animals were oriented backwards, most responses occurred at ±180° (where the stimuli appeared or disappeared at the edge of the monitor). These observations suggest that the PLEs have a wide visual field, from the end of the ALE range on one side (±60*°*), all the way around the back of the animal to the edge of the ALE range on the other side, for a total of ≈260*°*.

Regarding the validity of the scoring procedure for Experiment 2, as tested on the data from Experiment 1, we observed a significant preferential turning direction towards the stimulus position in the ALE treatments (GLMM post-hoc, Bonferroni corrected; estimated mean=16.306, SE=2.29, t=7.128, p<0.0001), while there was no such preference in PLE-only spiders (estimated mean=3.533, SE=2.34, t=1.511, p=0.5231). This second result was surprising to us, as the pivot clearly depends on the stimulus position (see Figure 2), and as such should be directed to the stimulus side, as we qualitatively observed. This may have been due to the lower response rate for the PLE treatment (ALE treatment response rate: 20.4% vs PLE treatment: 11.1%), which combined with the low sample size may have brought the effect under significance level. Moreover, in the PLE condition, especially for spiders facing frontally, the stimuli are for a long time outside the PLEs visual fields, accumulating a lot of motion independent from detection, contributing to decreasing the signal-to-noise ratio.

### 3.2 Experiment 2 – Eye-specific preference for biological motion

Results are summarized in Figure 3. Spiders in the ALE+PLE treatments showed no significant preference for either stimulus in the point-light-display pair (GLMM post-hoc, Bonferroni corrected; estimated mean=0.218, SE=1.44, t=0.151, p=1) nor in the shapes (silhouette vs ellipse) pair (estimated mean=-2.0592, SE=1.37, t=-1.506, p=0.7926). However, spiders in the ALE treatment showed a significant preference for the biological stimulus in the dots condition (estimated mean=7.5723, SE=2.52, t=3.003, p=0.0161) but no preference in the shapes condition (estimated mean=-1.9663, SE=2.32, t=-0.849, p=1). Lastly, spiders in the PLE treatment showed no significant preference for either stimulus in the dots condition (estimated mean=3.1869, SE=3.17, t=1.004, p=1) nor in the shapes condition (estimated mean=-0.0123, SE=2.55, t=-0.005, p=1).

**Figure 3.**
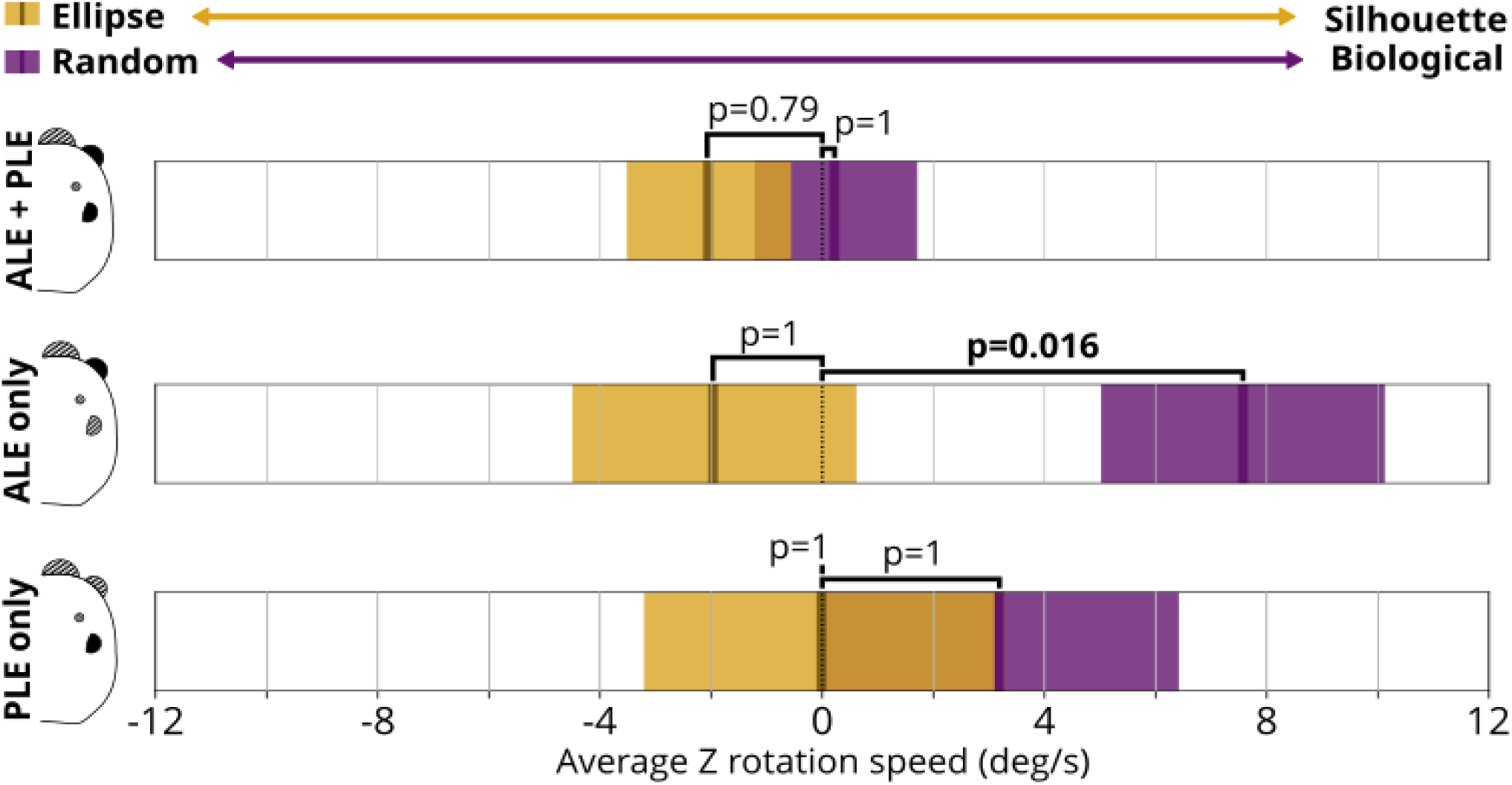
Results of experiment 2. on the x axis, the average rotational speed of the sphere Z axis is reported, in degrees per second. Negative numbers represent rotations consistent with the position of the non-biological option (Ellipse for gold bars, Random for purple), while positive values are rotations consistent with the position of the biological one (Silhouette for gold, Biological point light for purple). Dark bars represent the average, shaded region is standard error. We observed no rotational preference for either Ellipse or Silhouette in any condition. On the other hand, spiders were more prone to rotate towards the Biological option in respect to the Random one in the ALE only condition.

## 4 Discussion

In this paper, we tested the ability of jumping spiders to discriminate biological motion point-light displays from random ones. We did so under partial blindness conditions, to inquire about whether such discrimination ability is dedicated to a specific eye-pair.

We observed that the spiders were able to discriminate between random and biological displays with the ALEs unblocked, preferring the latter stimulus. They showed no preference when only the PLEs were unblocked. This confirms our initial hypothesis, as the biological motion recognition circuitry must be located in ALEs specific visual areas, attesting to the deep specialization of the modular visual system of jumping spiders.

We propose that biological motion detection is an ability dedicated to the ALEs likely in a very early area of their dedicated visual system and that it acts as a low level filter: neurons fire only when detecting local coherent motion (like biological point-light displays). If so, stimuli with fully incoherent local motion (i.e. random point-light-displays) would go completely undetected by the ALEs, with no neuronal firing carrying through to subsequent brain areas. As such, in the ALE treatment, the spider only detected the biological motion stimulus, and consistently pivoted towards it—as though the biological motion were the only stimulus presented. In contrast, the PLEs seem to act as simple motion detectors, with the relevant neural responses registering any translating stimulus. Thus, in the PLE treatment, as the two point-light displays translate across the screen at the same speed and with the same total motion amount, these two appeared identical to the spider, with equal information relayed to subsequent brain areas. This would cause no preference in the PLE treatment, as the two stimuli have no apparent difference.

Crucially, in our previous experiment (De Agrò et al., 2021) with all the eyes unblocked, the spiders turned more towards the random displays, not the biological ones. The opposite direction of preference in the ALE condition suggests multi-eye interaction to be particularly important in informing the spiders’ behavior. With all eyes available, a translating biological point-light display smoothly moves across the full spider field of view, starting from the PLEs and then passing over the ALE. As both eye-pairs are equally capable of detecting the stimulus, no mismatch is detected in the switch between the fields. If, however, the moving stimulus is a random point-light display, this will be detected by the PLEs, but will then unexpectedly disappear entering the ALEs. This sudden information mismatch may violate the spider’s “expectation”, causing an attention shift towards the disappeared object and as it will be recorded as a preference for the stimulus in our experiment (De Agrò et al., 2021). Note that with the term expectation we are not implying the necessity of a high-order representation of the object in the spider brain. This behavior can be instead produced with just three neuronal layers, fully fitting in the early areas of the spiders’ visual system. We provide a possible organization of such a system in Box 1. There is indeed a wealth of evidence that shows that spiders maximally produce pivots upon unexpected changes in stimuli behavior: obviously, when a stimulus first enters their field of view, but also when it stops, when it starts moving, when it leaves, or when it changes direction (De Agrò et al., 2021).

### Box 1 Computational model of jumping spiders’ pivoting behavior

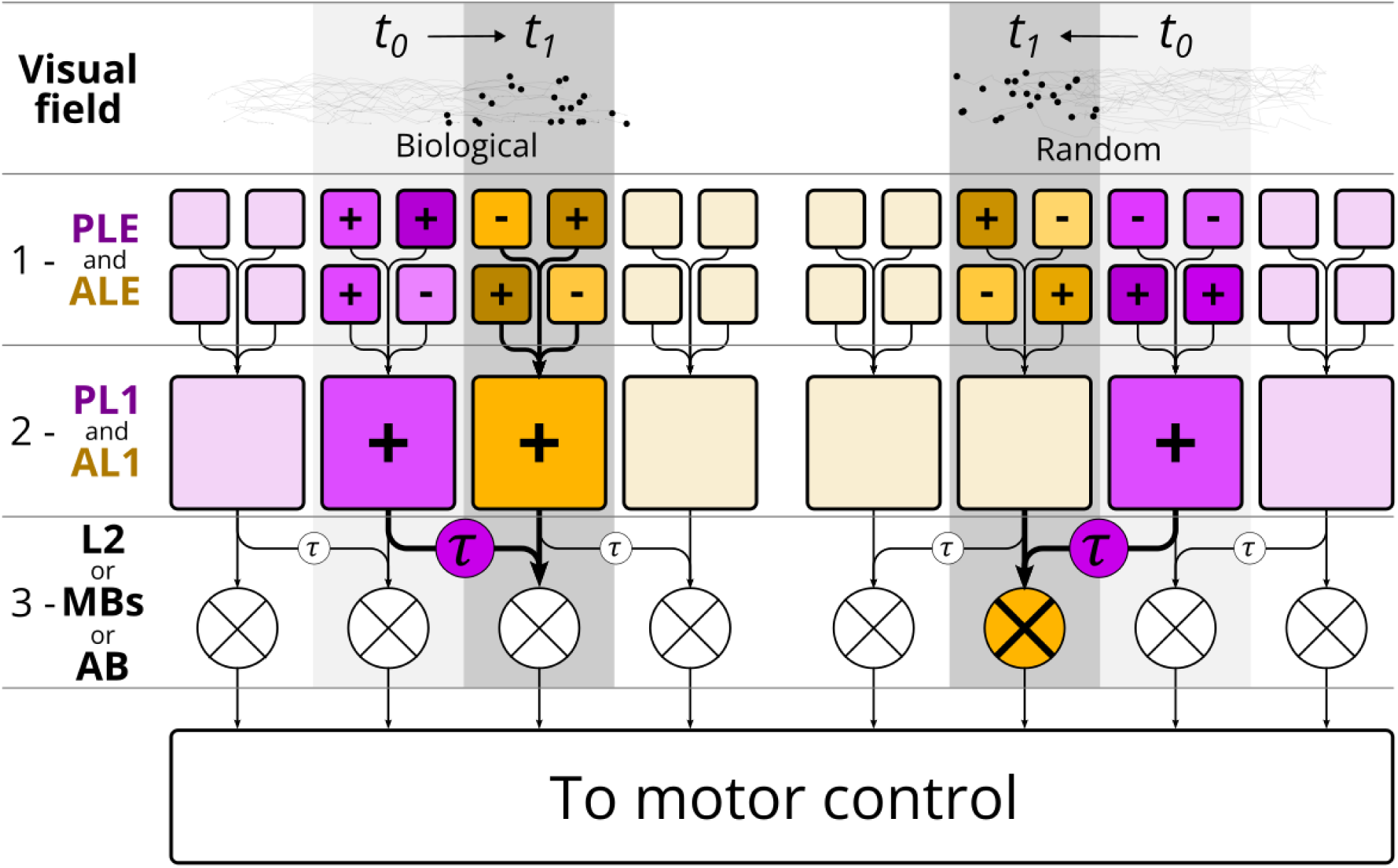

This hypothesized model attempts to account for the observed behavior of jumping spiders, switching preference for producing pivot towards biological vs random motion depending on the available eyes. The computational model is organized across 3 successive layers (numbered rows, 1-3). Layer 1 acts as the input layer, and it is composed of photosensitive cells; in the jumping spiders’ visual system this would correspond to the eyes (PLEs and ALEs specifically in the context of this experiment). Layer 2 cells collect input from multiple photoreceptors and extract motion types; in the jumping spiders’ visual system these would be located in the early, eye-specific visual areas (AL1 for ALEs, PL1 for PLEs. For a full description of the visual system organization, see Steinhoff et al., 2020). As per our hypothesis, these cells should be sensitive to specific types of motion: locally coherent for AL1, global direction for PL1. Layer 3 contains cells acting as exclusive or (XOR) gates, receiving direct input from a cell in layer 2, and delayed input from the neighboring ones; in the jumping spiders’ visual system, this would be located in a brain region receiving input from both AL1 and PL2 (e.g., the Mushroom Bodies, the Arcuate Body, L2). The direct and delayed connections presented here are a very similar system to the Reichardt detector (Haag et al., 2004; Reichardt, 1987), which describes how motion direction is encoded by the brain. The difference here is that rather than being directly connected to photoreceptors, the comparator and delayed connections are attached to a subsequent visual area. Purple boxes represent cells specific to PLEs and connected areas, while gold boxes represent cells specific to ALEs and connected areas. Brightly colored blocks represent active cells.

As in our experiment, let’s assume that across the visual field of the spider two stimuli are moving across, from the side towards the center. On the left, a biological point-light display, while on the right there is a random one. At time *t_0_* (light gray background), the two stimuli are moving across the edge of the PLEs field. The PLEs photoreceptors (layer 1) will react to changes in luminance on the visual field and send signals to the subsequent brain areas. As hypothesized, PLEs visual stream might be dedicated to global motion detection. As such, neurons in the dedicated brain region of these eyes (layer 2, PL1) will collect the pattern of activation of photoreceptors and react to the presence of the moving point-cloud. This will happen both for the biological and the random displays.

The stimuli will then continue to move, reaching the start of the ALEs field of view at time *t_1_*. ALEs photoreceptors (layer 1) will activate as well and send signal a to their dedicated brain region (layer 2, AL1). Crucially, AL1 neurons may be directly tuned for local motion coherency, and will react only for the biological motion pattern, but not for the random one. All neurons of AL1 and PL1 then will project to layer 3. Until a stimulus follows a predictive path, the XOR gates will not activate, as they will receive both the delayed signal of neurons attending to the stimulus position in *t_0_*, and the direct signal from the neurons attending the stimulus position in *t_1_*. However, since AL1 neurons do not fire for random motion, the signal will not carry over to the dedicated XOR gate, activating it due to a mismatch with the delayed connection and sending a signal to motor control. Pivot direction can be decided according to the relative activation of all the XOR neurons, turning towards the highest firing location. Note that this circuit can account also for stimuli appearing in the visual field, with the XOR gate receiving a signal from the direct connection, but no signal from the delayed one (as no activation occurred at the previous time step). The same is true for stimuli changing direction. Lastly, the system can also account for our PLEs only condition: both XOR gates connected to the two stimuli appearing locations will equally send signal to motor control, causing pivots to either direction randomly.

In the current experiment, however, the spiders did not show the same preference for random over biological displays in the ALEs+PLEs condition. We believe there are two possible explanations for the lack of preference. In the current experiments, the two computer monitors were placed at 65*°* to each other, with the two point light displays moving between ±90*°* and ±10*°*. This is in contrast with our previous study, where the stimuli moved between ±60*°* and ±5*°*. This means that in the current experiment the stimuli spent a much longer time passing across the PLEs field only (from ±90*°* to ±60*°*), leaving a long time for the spiders to pivot long before gaining information from the ALEs. In our previous experiment instead, the stimuli just barely appeared in the PLEs field, maximizing the importance of the PLEs/ALEs switch and driving the difference up. A second option may be linked to the unavailability of the AMEs. As previously stated, the pivots made towards random displays is fundamentally an information-seeking effort, directed towards a stimulus that violated expectations. Without AMEs, such a pivot would become redundant, as it would only bring the random stimulus frontally, fully in the ALEs field. Note that this behavior would still increase the information intake in the only ALEs and only PLEs conditions, as moving the detected target in the center of the ALEs field would at its minimum provide data about its distance (due to the fields overlap). Regardless, further inquiries would be needed to test which of these two explanations is the most appropriate, and replication is due to confirm this is not instead a type II error: spiders were able to discriminate the two stimuli, we just failed to observe it due to a low statistical power.

We found the spiders did not discriminate—that is, they showed no preference— between the silhouette and the ellipse. This was in line with our expectation, as the difference between the two is mostly shape-based and as such are more likely to be interpreted by AMEs. Indeed, even though the spider silhouette contains biological motion information, this is much less evident, as the absence of contrast and depth in the image hides the position of the leg joints, information that is instead enhanced in point-light-display stimuli. It has been argued that the spiders’ ALEs could also be capable of discriminating basic shapes, as their resolution should be sufficient (Goté et al., 2019). Behaviorally, however, this remains uncertain. Bruce et al. (2021) for example tested the effect of a distractor appearing in the ALEs field during AMEs scanning of a target stimulus. While the shape of the target influenced the probability of gaze shift, only the motion and not the shape of the distractor had an effect. Indeed, spatial acuity is not sufficient for shape recognition, as it requires dedicated circuitry that may be instead specific to AMEs, following our specialization hypothesis. Our experiment points in this direction, even though future studies will be needed to directly test the ability of ALEs regarding shape discrimination.

In this experiment, we demonstrated the presence of a deep specialization in the jumping spiders’ visual system. This specialization is much deeper than previously thought, with single eye-pairs fulfilling specific tasks. How every eye achieves recognition of motion as complex as biological displays still remain an open question. Here, we proposed the use of a cue like local coherency, but future studies will be needed to confirm this ALEs specialization, testing it directly with specifically designed stimuli. Moreover, we provided a testable hypothesis on how the interaction between the different eye-pairs may determine decision making, inflating the amount of information that each specialized pair can provide. We believe that moving complex computation upstream and exploiting mismatches is a unique solution to the problem of brain miniaturization and may be one solution for producing high performance with the limited resources.

## Supporting information

SI - analysis script and raw data

## 5 Acknowledgments

This research was funded by the Association for the Study of Animal Behaviour (ASAB) Research Grant to MDA (February 2021).

