## Supplementary material for "Eye-specific detection and a multi-eye integration model of biological motion perception": SI - analysis script and raw data: S1_Analysis.html

SM1 Analysis


### SM1 Analysis

Abstract

This supplement provides the entire R script and output of the
statistical analysis we performed and figures produced, in their
original form. It is presented in the spirit of open and transparent
science, but has not been carefully curated.

### Setup

#### Prepare R environment

```
library(readODS) #to read raw data
library(glmmTMB) #for mixed models
library(car) #for anova on mixed models
library(DHARMa) #for goodness of fit of the model
library(emmeans) #for post hoc
library(ggplot2) #to plot
library(reticulate)

use_python('/home/massimodeagro/anaconda3/envs/DataAnalysis/bin/python')
```

#### Prepare Python environment

```
import pandas as pd
import os
import matplotlib.pyplot as plt
import numpy as np
import seaborn as sns
from scipy.stats import gaussian_kde
```

#### Load data

I will load both Experiment 1 and 2 here

##### Experiment 1 - eye angle analysis

```
path = r.path
exp1 = pd.read_csv(path+"exp1.csv")
```

##### Experiment 1 - goodness of procedure control

##### Experiment 2 - Biological motion preference with specific eyes

```
main = pd.read_csv(path+"exp2_1.csv")
mainf = pd.read_csv(path+"exp2_2.csv")

exp2 = pd.concat([main, mainf])
```

### Analysis

#### Experiment 1

For experiment 1, we are not going to produce a proper statistical
analysis. The main questions are:

- what’s the pivot frequency for the different eyes and setups?
- at what stimulus angle do the pivot angle for the different
  eyes?

First, some general information

##### describing the data

```
nlevels(exp1$subj)
```

```
## [1] 31
```

```
summary(exp1$sex[!duplicated(exp1$subj)])
```

```
##  f  F  j  m 
## 13  1  8  9
```

```
summary(exp1$eyes[!duplicated(exp1$subj)])
```

```
## ALE PLE 
##  15  16
```

there are 31 subjects

14 females, 8 juveniles and 9 males

15 of ALE treatment, 16 for PLE

##### Pivot frequency

```
m1PivotFrequency <- glmmTMB(turn~eyes*screenangle*stimn+(trialn|subj), data=exp1, family = binomial)
simres <- simulateResiduals(m1PivotFrequency)
plot(simres)
```

```
Anova(m1PivotFrequency)
```

```
## Analysis of Deviance Table (Type II Wald chisquare tests)
## 
## Response: turn
##                          Chisq Df Pr(>Chisq)    
## eyes                    1.4895  1   0.222294    
## screenangle             7.2573  1   0.007061 ** 
## stimn                  18.0292  1  2.175e-05 ***
## eyes:screenangle        0.0009  1   0.976003    
## eyes:stimn              1.5758  1   0.209366    
## screenangle:stimn       3.7640  1   0.052368 .  
## eyes:screenangle:stimn  4.1831  1   0.040829 *  
## ---
## Signif. codes:  0 '***' 0.001 '**' 0.01 '*' 0.05 '.' 0.1 ' ' 1
```

Indeed, there seems to be a difference in reaction rate for eyes and
for the angle. Not of the interaction, but we would not have expected to
find it.

```
et <- emtrends(m1PivotFrequency, ~eyes*screenangle, var='stimn', type='response')
et
```

```
##  eyes screenangle stimn.trend     SE  df asymp.LCL asymp.UCL
##  ALE  65             -0.05296 0.0168 Inf   -0.0859   -0.0201
##  PLE  65             -0.00237 0.0158 Inf   -0.0334    0.0286
##  ALE  120            -0.05540 0.0212 Inf   -0.0969   -0.0139
##  PLE  120            -0.08792 0.0259 Inf   -0.1387   -0.0371
## 
## Confidence level used: 0.95
```

```
test(et, adjust='bonferroni')
```

```
##  eyes screenangle stimn.trend     SE  df z.ratio p.value
##  ALE  65             -0.05296 0.0168 Inf  -3.156  0.0064
##  PLE  65             -0.00237 0.0158 Inf  -0.150  1.0000
##  ALE  120            -0.05540 0.0212 Inf  -2.619  0.0352
##  PLE  120            -0.08792 0.0259 Inf  -3.391  0.0028
## 
## P value adjustment: bonferroni method for 4 tests
```

```
e <- emmeans(m1PivotFrequency, ~eyes*screenangle, type='response')
```

```
## NOTE: Results may be misleading due to involvement in interactions
```

```
e
```

```
##  eyes screenangle   prob     SE  df asymp.LCL asymp.UCL
##  ALE  65          0.1820 0.0663 Inf    0.0851    0.3475
##  PLE  65          0.1073 0.0381 Inf    0.0522    0.2077
##  ALE  120         0.0934 0.0418 Inf    0.0377    0.2133
##  PLE  120         0.0421 0.0187 Inf    0.0174    0.0982
## 
## Confidence level used: 0.95 
## Intervals are back-transformed from the logit scale
```

```
contrast(e, list('ALE vs PLE' = c(0.5, -0.5, 0.5, -0.5),
                 '65 vs 120' = c(0.5, 0.5, -0.5, -0.5)), adjust = 'bonferroni')
```

```
##  contrast   odds.ratio    SE  df null z.ratio p.value
##  ALE vs PLE       2.08 1.207 Inf    1   1.269  0.4090
##  65 vs 120        2.43 0.713 Inf    1   3.028  0.0049
## 
## P value adjustment: bonferroni method for 2 tests 
## Tests are performed on the log odds ratio scale
```

As a first evidence, we can observe a decrease in response rate
across every trial. This has to be expected.

Moreover, we found that spiders respond to stimuli more often in the
ALE treatment in respect to the PLE treatment. The different is not
statistically significant probably for the low n, but ALE response is
almost double. This is consistent with evidences identifying the ALE as
the primary control eyes for gaze direction of AME.

Lastly, Response rate is higher for the 65deg condition over the
120deg one. We will use this for the next experiment.

##### pivoting angle

let’s make density plots of the response position for the different
eyes. I will also divide for the PLEs the forward and backward spider
positioning, as the visible angles change drastically

```
ALE = exp1[exp1['eyes']=='ALE']
PLE = exp1[exp1['eyes']=='PLE']

PLErev = PLE[PLE['reversed']==1]
PLEstraight = PLE[PLE['reversed']==0]

plt.rcParams['font.size'] = '16'

fig, ax = plt.subplots(1,2, subplot_kw={'projection': 'polar'}, )

ax[0].hist(np.deg2rad(ALE['angposabs']), bins=15, density=True, alpha=0.8)
```

```
## (array([0.16477838, 0.65911352, 0.60418739, 2.08719282, 1.53793155,
##        0.27463063, 0.43940901, 0.49433514, 0.43940901, 0.43940901,
##        0.21970451, 0.21970451, 0.05492613, 0.21970451, 0.21970451]), array([0.42844076, 0.55229295, 0.67614514, 0.79999733, 0.92384952,
##        1.04770171, 1.1715539 , 1.29540609, 1.41925828, 1.54311047,
##        1.66696266, 1.79081485, 1.91466704, 2.03851923, 2.16237142,
##        2.28622361]), <BarContainer object of 15 artists>)
```

```
ax[0].set_thetamin(0)
ax[0].set_thetamax(180)
ax[0].set_theta_zero_location("N")
ax[0].set_rlim(-1,2.1)
```

```
## (-1.0, 2.1)
```

```
ax[0].set_rlabel_position(-22.5)  # Move radial labels away from plotted line
ax[0].grid(True)

ax[1].hist(np.deg2rad(-PLEstraight['angposabs']), bins=15, density=True, alpha=0.8)
```

```
## (array([0.73818396, 0.36909198, 0.92272995, 0.        , 0.36909198,
##        0.92272995, 0.18454599, 0.18454599, 0.        , 1.66091391,
##        1.10727594, 0.18454599, 0.36909198, 0.55363797, 0.36909198]), array([-2.28622361, -2.16020725, -2.03419088, -1.90817452, -1.78215816,
##        -1.6561418 , -1.53012544, -1.40410908, -1.27809272, -1.15207636,
##        -1.02606   , -0.90004364, -0.77402728, -0.64801092, -0.52199456,
##        -0.3959782 ]), <BarContainer object of 15 artists>)
```

```
ax[1].hist(np.deg2rad(-PLErev['angposabs']), bins=15, density=True, alpha=0.8)
```

```
## (array([1.4677289 , 0.        , 0.20967556, 0.41935111, 0.41935111,
##        0.41935111, 0.52418889, 0.20967556, 0.83870223, 0.41935111,
##        0.31451334, 0.41935111, 0.31451334, 0.20967556, 0.52418889]), array([-3.13569086, -2.98665107, -2.83761129, -2.6885715 , -2.53953172,
##        -2.39049193, -2.24145215, -2.09241236, -1.94337258, -1.79433279,
##        -1.64529301, -1.49625322, -1.34721343, -1.19817365, -1.04913386,
##        -0.90009408]), <BarContainer object of 15 artists>)
```

```
ax[1].set_thetamin(0)
ax[1].set_thetamax(-180)
ax[1].set_theta_zero_location("N")
ax[1].set_rlim(-1,2.1)
```

```
## (-1.0, 2.1)
```

```
ax[1].set_rlabel_position(-22.5)  # Move radial labels away from plotted line
ax[1].grid(True)

fig.show()
```

For some reason that escapes me, the plot is not getting on the html.
This is not crucial as this plot is being included in the main paper, so
we will not waste too much time fixing it.

The response for ALE is very clear: a peak is appreciable at the 50°
mark. This is sufficient to identify the ALE visual field as starting at
+-50°

For the PLE the story is more complex. When the spider is facing the
screens, and as such available angles are between +-120°, a clear peak
can be seen at 60°. this could very well be the start of the PLE visual
field. When the spider is facing backward, and as such available angles
are between +-60° (passing across +-180° and excluding frontal ones) a
peak at 180° is appreciable. This is when the stimuli appear from the
contact corner of the two monitors. Our interpretation is that actually
the spider can see across the whole visual field span from 60° to
-60°.

Regardless, for the current experiment this is not crucial. We can
select to show the stimuli of our main experiment across the angles 30
to 70, to cover the area in which both eyes are sensitive.

##### Control for scoring of next experiment

As in De Agrò et al 2021, the scoring for experiment 2 will be
performed by collecting all peaks in the signal, setting as positive the
ones according with the position of the bio stim and as negative the
ones according with the position of the random stim. Averaging out, a
positive value will correspond to a preference for bio, negative with a
preference for random, 0 for no preference. We will use the same scoring
procedure here, where there is only one stimulus. The result should turn
out positive, as most if not all rotations should happen towards the
stimulus.

```
hist(exp1C$dirval_deg_s, breaks=100)
```

```
mcontrol <- glmmTMB(dirval_deg_s~stiminfocus*eyes + (1|subj), #I am dropping trialn as random slope as the model fails to converge. Should not be a problem as all trials are the same
                 data=exp1C, family= gaussian(),
                 control=glmmTMBControl(optCtrl = list(iter.max = 300000, eval.max = 400000)))

simres <- simulateResiduals(mcontrol)
plot(simres)
```

Even though Dharma gives error, it is clearly a gaussian from the
histogram

```
Anova(mcontrol)
```

```
## Analysis of Deviance Table (Type II Wald chisquare tests)
## 
## Response: dirval_deg_s
##                    Chisq Df Pr(>Chisq)    
## stiminfocus      37.4679  1  9.293e-10 ***
## eyes              0.0002  1     0.9889    
## stiminfocus:eyes 25.8300  1  3.729e-07 ***
## ---
## Signif. codes:  0 '***' 0.001 '**' 0.01 '*' 0.05 '.' 0.1 ' ' 1
```

```
e <- emmeans(mcontrol, ~eyes*stiminfocus, type='response')
e
```

```
##  eyes stiminfocus emmean   SE    df lower.CL upper.CL
##  ALE            0  0.373 1.56 10782   -2.689     3.44
##  PLE            0  2.207 1.45 10782   -0.642     5.06
##  ALE            1 16.306 2.29 10782   11.822    20.79
##  PLE            1  3.533 2.34 10782   -1.050     8.12
## 
## Confidence level used: 0.95
```

```
test(e, adjust='bonferroni')
```

```
##  eyes stiminfocus emmean   SE    df t.ratio p.value
##  ALE            0  0.373 1.56 10782   0.239  1.0000
##  PLE            0  2.207 1.45 10782   1.519  0.5156
##  ALE            1 16.306 2.29 10782   7.128  <.0001
##  PLE            1  3.533 2.34 10782   1.511  0.5231
## 
## P value adjustment: bonferroni method for 4 tests
```

The result for ALE is what we would expect. For PLE the thing is more
complicated. The value is positive, but not significant. This could be
due to the lower amount of “meaningful” responses, as shown in the
binomial check above. This will make the relative amount of noise
increase, other than decreasing the statistical power. This poses an
interpretation challange to the main experiment data: if with one
stimulus no preference shows up, we cannot expect a preference to appear
in the main experiment.

That said, the angle analysis shows clear peaks at 120 and 60, as
shown above. This must mean that the spiders are indeed turning at
meaningful moments, and so our measure must be consistent with real
pivoting.

#### Experiment 2

##### describing the data

```
nlevels(exp2$subj)
```

```
## [1] 179
```

```
summary(exp2$sex[!duplicated(exp2$subj)])
```

```
##  f  j  m 
## 89 75 15
```

```
summary(exp2$eyes[!duplicated(exp2$subj)])
```

```
##     ALE Control     PLE 
##      58      61      60
```

##### Preliminary analysis

Before proceeding with the main analysis, we will perform some checks
on the data

###### where do rotations happen?

In De Agrò et al. 2021, rotations were consistently associated with
the presence of the stimulus, with a correlation between rotational
magnitude and stimulus angular position. We did also observe that when
the stimuli started after being static, there was an increase in
response. In this experiment only one stop-start point was set, which
may change our observation. Also, the fact that stimuli come from the
center of the FOV and go out in some cases may trigger more rotation for
some angles, differently from what previously observed. Lastly,
depending on the eye pair, stimuli at the same visual angle may be
entering or leaving the visual field, changing completely the expected
reaction. Let’s observe the data to see if such differences are
relevant.

```
hist(exp2$absval_deg_s, breaks=50)
```

```
mspeed <- glmmTMB(absval_deg_s~stimpos*stimdir*eyes*cond + (cond|subj),
                data = exp2, family = Gamma(link='log'),
                control=glmmTMBControl(optCtrl = list(iter.max = 30000, eval.max = 40000)))

simres <- simulateResiduals(mspeed)
plot(simres, factor=TRUE)
```

```
## Warning in plot.window(...): parametro grafico "factor" non valido
```

```
## Warning in plot.xy(xy, type, ...): parametro grafico "factor" non valido
```

```
## Warning in title(...): parametro grafico "factor" non valido
```

the model is not perfect, but good enough for the fit looking at the
histogram too.

```
summary(mspeed)
```

```
##  Family: Gamma  ( log )
## Formula:          
## absval_deg_s ~ stimpos * stimdir * eyes * cond + (cond | subj)
## Data: exp2
## 
##      AIC      BIC   logLik deviance df.resid 
## 186509.6 186730.3 -93226.8 186453.6    19569 
## 
## Random effects:
## 
## Conditional model:
##  Groups Name             Variance Std.Dev. Corr  
##  subj   (Intercept)      0.1173   0.3425         
##         condsilh-ellipse 0.1188   0.3447   -0.54 
## Number of obs: 19597, groups:  subj, 178
## 
## Dispersion estimate for Gamma family (sigma^2): 0.775 
## 
## Conditional model:
##                                                       Estimate Std. Error
## (Intercept)                                          3.8611587  0.0746440
## stimpos                                             -0.0020059  0.0008714
## stimdiroutward                                      -0.2292200  0.0737812
## eyesControl                                         -0.2386393  0.1122125
## eyesPLE                                             -0.2936575  0.1086414
## condsilh-ellipse                                    -0.1130079  0.0887855
## stimpos:stimdiroutward                               0.0018978  0.0011782
## stimpos:eyesControl                                  0.0025569  0.0013118
## stimpos:eyesPLE                                      0.0021170  0.0012679
## stimdiroutward:eyesControl                           0.1886819  0.1107014
## stimdiroutward:eyesPLE                               0.2458572  0.1098141
## stimpos:condsilh-ellipse                             0.0002836  0.0011431
## stimdiroutward:condsilh-ellipse                      0.0477855  0.0990363
## eyesControl:condsilh-ellipse                         0.0609780  0.1380445
## eyesPLE:condsilh-ellipse                             0.2975437  0.1311257
## stimpos:stimdiroutward:eyesControl                  -0.0005157  0.0017802
## stimpos:stimdiroutward:eyesPLE                      -0.0004594  0.0017889
## stimpos:stimdiroutward:condsilh-ellipse              0.0007240  0.0016391
## stimpos:eyesControl:condsilh-ellipse                 0.0020176  0.0017790
## stimpos:eyesPLE:condsilh-ellipse                    -0.0004580  0.0016980
## stimdiroutward:eyesControl:condsilh-ellipse         -0.0335840  0.1525444
## stimdiroutward:eyesPLE:condsilh-ellipse             -0.2935646  0.1483306
## stimpos:stimdiroutward:eyesControl:condsilh-ellipse -0.0007274  0.0025040
## stimpos:stimdiroutward:eyesPLE:condsilh-ellipse      0.0029401  0.0024920
##                                                     z value Pr(>|z|)    
## (Intercept)                                           51.73  < 2e-16 ***
## stimpos                                               -2.30  0.02134 *  
## stimdiroutward                                        -3.11  0.00189 ** 
## eyesControl                                           -2.13  0.03345 *  
## eyesPLE                                               -2.70  0.00687 ** 
## condsilh-ellipse                                      -1.27  0.20308    
## stimpos:stimdiroutward                                 1.61  0.10722    
## stimpos:eyesControl                                    1.95  0.05127 .  
## stimpos:eyesPLE                                        1.67  0.09498 .  
## stimdiroutward:eyesControl                             1.70  0.08830 .  
## stimdiroutward:eyesPLE                                 2.24  0.02517 *  
## stimpos:condsilh-ellipse                               0.25  0.80406    
## stimdiroutward:condsilh-ellipse                        0.48  0.62945    
## eyesControl:condsilh-ellipse                           0.44  0.65869    
## eyesPLE:condsilh-ellipse                               2.27  0.02326 *  
## stimpos:stimdiroutward:eyesControl                    -0.29  0.77206    
## stimpos:stimdiroutward:eyesPLE                        -0.26  0.79732    
## stimpos:stimdiroutward:condsilh-ellipse                0.44  0.65868    
## stimpos:eyesControl:condsilh-ellipse                   1.13  0.25677    
## stimpos:eyesPLE:condsilh-ellipse                      -0.27  0.78735    
## stimdiroutward:eyesControl:condsilh-ellipse           -0.22  0.82575    
## stimdiroutward:eyesPLE:condsilh-ellipse               -1.98  0.04780 *  
## stimpos:stimdiroutward:eyesControl:condsilh-ellipse   -0.29  0.77145    
## stimpos:stimdiroutward:eyesPLE:condsilh-ellipse        1.18  0.23807    
## ---
## Signif. codes:  0 '***' 0.001 '**' 0.01 '*' 0.05 '.' 0.1 ' ' 1
```

```
Anova(mspeed)
```

```
## Warning in printHypothesis(L, rhs, names(b)): one or more coefficients in the hypothesis include
##      arithmetic operators in their names;
##   the printed representation of the hypothesis will be omitted

## Warning in printHypothesis(L, rhs, names(b)): one or more coefficients in the hypothesis include
##      arithmetic operators in their names;
##   the printed representation of the hypothesis will be omitted

## Warning in printHypothesis(L, rhs, names(b)): one or more coefficients in the hypothesis include
##      arithmetic operators in their names;
##   the printed representation of the hypothesis will be omitted

## Warning in printHypothesis(L, rhs, names(b)): one or more coefficients in the hypothesis include
##      arithmetic operators in their names;
##   the printed representation of the hypothesis will be omitted

## Warning in printHypothesis(L, rhs, names(b)): one or more coefficients in the hypothesis include
##      arithmetic operators in their names;
##   the printed representation of the hypothesis will be omitted

## Warning in printHypothesis(L, rhs, names(b)): one or more coefficients in the hypothesis include
##      arithmetic operators in their names;
##   the printed representation of the hypothesis will be omitted

## Warning in printHypothesis(L, rhs, names(b)): one or more coefficients in the hypothesis include
##      arithmetic operators in their names;
##   the printed representation of the hypothesis will be omitted

## Warning in printHypothesis(L, rhs, names(b)): one or more coefficients in the hypothesis include
##      arithmetic operators in their names;
##   the printed representation of the hypothesis will be omitted

## Warning in printHypothesis(L, rhs, names(b)): one or more coefficients in the hypothesis include
##      arithmetic operators in their names;
##   the printed representation of the hypothesis will be omitted

## Warning in printHypothesis(L, rhs, names(b)): one or more coefficients in the hypothesis include
##      arithmetic operators in their names;
##   the printed representation of the hypothesis will be omitted

## Warning in printHypothesis(L, rhs, names(b)): one or more coefficients in the hypothesis include
##      arithmetic operators in their names;
##   the printed representation of the hypothesis will be omitted

## Warning in printHypothesis(L, rhs, names(b)): one or more coefficients in the hypothesis include
##      arithmetic operators in their names;
##   the printed representation of the hypothesis will be omitted

## Warning in printHypothesis(L, rhs, names(b)): one or more coefficients in the hypothesis include
##      arithmetic operators in their names;
##   the printed representation of the hypothesis will be omitted

## Warning in printHypothesis(L, rhs, names(b)): one or more coefficients in the hypothesis include
##      arithmetic operators in their names;
##   the printed representation of the hypothesis will be omitted

## Warning in printHypothesis(L, rhs, names(b)): one or more coefficients in the hypothesis include
##      arithmetic operators in their names;
##   the printed representation of the hypothesis will be omitted
```

```
## Analysis of Deviance Table (Type II Wald chisquare tests)
## 
## Response: absval_deg_s
##                             Chisq Df Pr(>Chisq)    
## stimpos                   11.6974  1  0.0006259 ***
## stimdir                    0.0122  1  0.9119356    
## eyes                       0.6911  2  0.7078210    
## cond                       1.9607  1  0.1614415    
## stimpos:stimdir           20.2259  1  6.881e-06 ***
## stimpos:eyes              28.3204  2  7.085e-07 ***
## stimdir:eyes              21.8110  2  1.836e-05 ***
## stimpos:cond               7.3498  1  0.0067071 ** 
## stimdir:cond               0.4945  1  0.4819481    
## eyes:cond                  6.7100  2  0.0349089 *  
## stimpos:stimdir:eyes       2.0772  2  0.3539581    
## stimpos:stimdir:cond       1.8318  1  0.1759203    
## stimpos:eyes:cond          1.7091  2  0.4254845    
## stimdir:eyes:cond          4.3352  2  0.1144504    
## stimpos:stimdir:eyes:cond  2.1746  2  0.3371315    
## ---
## Signif. codes:  0 '***' 0.001 '**' 0.01 '*' 0.05 '.' 0.1 ' ' 1
```

Indeed, spider rotation is correlated with stimulus position, but
there is a big correlation with stimulus direction (inward or outward).
moreover there are interactions with uncovered eyes and condition, as we
expected. post-hoc follows for stimpos and stimdir

```
t <- emtrends(mspeed, ~stimdir*eyes, var='stimpos')
```

```
## NOTE: Results may be misleading due to involvement in interactions
```

```
test(t, adjust="bonferroni")
```

```
##  stimdir eyes    stimpos.trend       SE  df z.ratio p.value
##  inward  ALE         -1.86e-03 0.000572 Inf  -3.259  0.0067
##  outward ALE          3.96e-04 0.000586 Inf   0.675  1.0000
##  inward  Control      1.70e-03 0.000682 Inf   2.496  0.0754
##  outward Control      3.08e-03 0.000648 Inf   4.754  <.0001
##  inward  PLE          2.38e-05 0.000628 Inf   0.038  1.0000
##  outward PLE          3.29e-03 0.000697 Inf   4.728  <.0001
## 
## Results are averaged over the levels of: cond 
## P value adjustment: bonferroni method for 6 tests
```

```
fig, axs = plt.subplots(2,3)

focus= exp2[exp2['eyes']=="Control"]
axs[0,0].hist(focus[focus['stimdir']=="inward"]['timefromonset'].values, 100, density=True, facecolor='#593d9c', alpha=0.5)
```

```
## (array([0.05212382, 0.04397948, 0.04072174, 0.04343652, 0.04778017,
##        0.04397948, 0.04832313, 0.04778017, 0.04506539, 0.04072174,
##        0.03963582, 0.0336633 , 0.03203443, 0.02877669, 0.0336633 ,
##        0.03149148, 0.03692104, 0.03203443, 0.04343652, 0.04506539,
##        0.04669426, 0.03800696, 0.04669426, 0.04289356, 0.03800696,
##        0.04778017, 0.05538156, 0.05701043, 0.05918226, 0.04289356,
##        0.03800696, 0.03583513, 0.03474922, 0.02877669, 0.01846052,
##        0.0211753 , 0.02226122, 0.02769078, 0.02443304, 0.024976  ,
##        0.03149148, 0.02877669, 0.03094852, 0.02551896, 0.03257739,
##        0.0336633 , 0.03203443, 0.02769078, 0.0336633 , 0.03637809,
##        0.03149148, 0.03800696, 0.03529217, 0.0336633 , 0.03909287,
##        0.03963582, 0.04072174, 0.03692104, 0.03692104, 0.03637809,
##        0.037464  , 0.03637809, 0.04560835, 0.04669426, 0.04832313,
##        0.04560835, 0.04778017, 0.04669426, 0.04452243, 0.04017878,
##        0.04723722, 0.0461513 , 0.04669426, 0.04289356, 0.04723722,
##        0.05212382, 0.04397948, 0.049952  , 0.05158087, 0.04397948,
##        0.04778017, 0.04832313, 0.05158087, 0.04940904, 0.04832313,
##        0.049952  , 0.04235061, 0.05158087, 0.05158087, 0.05049496,
##        0.05429565, 0.05266678, 0.04560835, 0.04126469, 0.04940904,
##        0.05049496, 0.04017878, 0.0461513 , 0.04723722, 0.00597252]), array([2.47859955e-03, 2.47688046e-01, 4.92897491e-01, 7.38106937e-01,
##        9.83316383e-01, 1.22852583e+00, 1.47373528e+00, 1.71894472e+00,
##        1.96415417e+00, 2.20936361e+00, 2.45457306e+00, 2.69978251e+00,
##        2.94499195e+00, 3.19020140e+00, 3.43541084e+00, 3.68062029e+00,
##        3.92582973e+00, 4.17103918e+00, 4.41624863e+00, 4.66145807e+00,
##        4.90666752e+00, 5.15187696e+00, 5.39708641e+00, 5.64229586e+00,
##        5.88750530e+00, 6.13271475e+00, 6.37792419e+00, 6.62313364e+00,
##        6.86834309e+00, 7.11355253e+00, 7.35876198e+00, 7.60397142e+00,
##        7.84918087e+00, 8.09439032e+00, 8.33959976e+00, 8.58480921e+00,
##        8.83001865e+00, 9.07522810e+00, 9.32043755e+00, 9.56564699e+00,
##        9.81085644e+00, 1.00560659e+01, 1.03012753e+01, 1.05464848e+01,
##        1.07916942e+01, 1.10369037e+01, 1.12821131e+01, 1.15273226e+01,
##        1.17725320e+01, 1.20177415e+01, 1.22629509e+01, 1.25081603e+01,
##        1.27533698e+01, 1.29985792e+01, 1.32437887e+01, 1.34889981e+01,
##        1.37342076e+01, 1.39794170e+01, 1.42246265e+01, 1.44698359e+01,
##        1.47150454e+01, 1.49602548e+01, 1.52054642e+01, 1.54506737e+01,
##        1.56958831e+01, 1.59410926e+01, 1.61863020e+01, 1.64315115e+01,
##        1.66767209e+01, 1.69219304e+01, 1.71671398e+01, 1.74123493e+01,
##        1.76575587e+01, 1.79027682e+01, 1.81479776e+01, 1.83931870e+01,
##        1.86383965e+01, 1.88836059e+01, 1.91288154e+01, 1.93740248e+01,
##        1.96192343e+01, 1.98644437e+01, 2.01096532e+01, 2.03548626e+01,
##        2.06000721e+01, 2.08452815e+01, 2.10904910e+01, 2.13357004e+01,
##        2.15809098e+01, 2.18261193e+01, 2.20713287e+01, 2.23165382e+01,
##        2.25617476e+01, 2.28069571e+01, 2.30521665e+01, 2.32973760e+01,
##        2.35425854e+01, 2.37877949e+01, 2.40330043e+01, 2.42782137e+01,
##        2.45234232e+01]), <BarContainer object of 100 artists>)
```

```
axs[0,0].set_title("Control Inward")
axs[1,0].hist(focus[focus['stimdir']=="outward"]['timefromonset'].values, 100, density=True, facecolor='#593d9c', alpha=0.5)
```

```
## (array([0.04361958, 0.05045157, 0.04940049, 0.05307925, 0.04309405,
##        0.04992603, 0.06569215, 0.06253892, 0.0509771 , 0.04151744,
##        0.0409919 , 0.03731314, 0.04046636, 0.03468545, 0.03573653,
##        0.02732793, 0.02785347, 0.02995562, 0.02890454, 0.03888975,
##        0.04256851, 0.04204297, 0.03888975, 0.03731314, 0.0367876 ,
##        0.0451962 , 0.05045157, 0.05255372, 0.04046636, 0.04151744,
##        0.02943008, 0.01944488, 0.02207256, 0.02943008, 0.02154702,
##        0.03363438, 0.02364917, 0.03048116, 0.0325833 , 0.03363438,
##        0.03205777, 0.02417471, 0.03415992, 0.03415992, 0.03836421,
##        0.03888975, 0.04414512, 0.03468545, 0.04046636, 0.03626206,
##        0.03521099, 0.04046636, 0.03836421, 0.04361958, 0.03731314,
##        0.03626206, 0.03468545, 0.03994082, 0.04361958, 0.04204297,
##        0.0367876 , 0.04309405, 0.04361958, 0.04414512, 0.03836421,
##        0.04046636, 0.03888975, 0.04624727, 0.03626206, 0.04309405,
##        0.04046636, 0.04151744, 0.04467066, 0.04940049, 0.04467066,
##        0.05570694, 0.05833463, 0.0409919 , 0.04046636, 0.04256851,
##        0.04046636, 0.04887496, 0.05150264, 0.05413033, 0.04361958,
##        0.04256851, 0.05465587, 0.04729834, 0.04309405, 0.04992603,
##        0.04834942, 0.0451962 , 0.05202818, 0.04572173, 0.04729834,
##        0.03836421, 0.05413033, 0.04151744, 0.04992603, 0.0141895 ]), array([8.36849213e-05, 2.44724338e-01, 4.89364991e-01, 7.34005644e-01,
##        9.78646297e-01, 1.22328695e+00, 1.46792760e+00, 1.71256826e+00,
##        1.95720891e+00, 2.20184956e+00, 2.44649022e+00, 2.69113087e+00,
##        2.93577152e+00, 3.18041218e+00, 3.42505283e+00, 3.66969348e+00,
##        3.91433414e+00, 4.15897479e+00, 4.40361544e+00, 4.64825609e+00,
##        4.89289675e+00, 5.13753740e+00, 5.38217805e+00, 5.62681871e+00,
##        5.87145936e+00, 6.11610001e+00, 6.36074067e+00, 6.60538132e+00,
##        6.85002197e+00, 7.09466263e+00, 7.33930328e+00, 7.58394393e+00,
##        7.82858459e+00, 8.07322524e+00, 8.31786589e+00, 8.56250654e+00,
##        8.80714720e+00, 9.05178785e+00, 9.29642850e+00, 9.54106916e+00,
##        9.78570981e+00, 1.00303505e+01, 1.02749911e+01, 1.05196318e+01,
##        1.07642724e+01, 1.10089131e+01, 1.12535537e+01, 1.14981944e+01,
##        1.17428350e+01, 1.19874757e+01, 1.22321163e+01, 1.24767570e+01,
##        1.27213976e+01, 1.29660383e+01, 1.32106790e+01, 1.34553196e+01,
##        1.36999603e+01, 1.39446009e+01, 1.41892416e+01, 1.44338822e+01,
##        1.46785229e+01, 1.49231635e+01, 1.51678042e+01, 1.54124448e+01,
##        1.56570855e+01, 1.59017261e+01, 1.61463668e+01, 1.63910074e+01,
##        1.66356481e+01, 1.68802888e+01, 1.71249294e+01, 1.73695701e+01,
##        1.76142107e+01, 1.78588514e+01, 1.81034920e+01, 1.83481327e+01,
##        1.85927733e+01, 1.88374140e+01, 1.90820546e+01, 1.93266953e+01,
##        1.95713359e+01, 1.98159766e+01, 2.00606172e+01, 2.03052579e+01,
##        2.05498985e+01, 2.07945392e+01, 2.10391799e+01, 2.12838205e+01,
##        2.15284612e+01, 2.17731018e+01, 2.20177425e+01, 2.22623831e+01,
##        2.25070238e+01, 2.27516644e+01, 2.29963051e+01, 2.32409457e+01,
##        2.34855864e+01, 2.37302270e+01, 2.39748677e+01, 2.42195083e+01,
##        2.44641490e+01]), <BarContainer object of 100 artists>)
```

```
axs[1,0].set_title("Control Outward")

focus= exp2[exp2['eyes']=="ALE"]
axs[0,1].hist(focus[focus['stimdir']=="inward"]['timefromonset'].values, 100, density=True, facecolor='#593d9c', alpha=0.5)
```

```
## (array([0.05303619, 0.04825097, 0.04545959, 0.04306698, 0.05503003,
##        0.05343496, 0.05144112, 0.04984604, 0.04585836, 0.04187068,
##        0.04386452, 0.04067437, 0.05223865, 0.06220786, 0.06061279,
##        0.05981525, 0.05822018, 0.05861895, 0.0566251 , 0.04386452,
##        0.04864974, 0.04745343, 0.06779062, 0.09012164, 0.09690071,
##        0.09769824, 0.07775983, 0.07018323, 0.06021402, 0.03748422,
##        0.02472364, 0.02193226, 0.01914088, 0.01993842, 0.01993842,
##        0.01993842, 0.01834334, 0.0163495 , 0.01754581, 0.02352733,
##        0.01874211, 0.01993842, 0.01914088, 0.02352733, 0.01834334,
##        0.02591994, 0.02512241, 0.02631871, 0.02711625, 0.02591994,
##        0.02711625, 0.03030639, 0.02711625, 0.03230024, 0.03110393,
##        0.03509161, 0.02990763, 0.03708546, 0.03110393, 0.03828176,
##        0.03230024, 0.03708546, 0.03748422, 0.03429408, 0.03509161,
##        0.04226944, 0.04067437, 0.0402756 , 0.032699  , 0.03868053,
##        0.04426329, 0.04107314, 0.03708546, 0.04226944, 0.0478522 ,
##        0.03828176, 0.03987683, 0.04346575, 0.04585836, 0.04386452,
##        0.0466559 , 0.04426329, 0.04825097, 0.04386452, 0.04226944,
##        0.04466205, 0.03110393, 0.00079754, 0.00039877, 0.00039877,
##        0.        , 0.00039877, 0.        , 0.00039877, 0.        ,
##        0.00039877, 0.00039877, 0.00039877, 0.00039877, 0.00079754]), array([2.43282318e-03, 2.82562587e-01, 5.62692351e-01, 8.42822115e-01,
##        1.12295188e+00, 1.40308164e+00, 1.68321141e+00, 1.96334117e+00,
##        2.24347094e+00, 2.52360070e+00, 2.80373046e+00, 3.08386023e+00,
##        3.36398999e+00, 3.64411976e+00, 3.92424952e+00, 4.20437928e+00,
##        4.48450905e+00, 4.76463881e+00, 5.04476858e+00, 5.32489834e+00,
##        5.60502810e+00, 5.88515787e+00, 6.16528763e+00, 6.44541740e+00,
##        6.72554716e+00, 7.00567693e+00, 7.28580669e+00, 7.56593645e+00,
##        7.84606622e+00, 8.12619598e+00, 8.40632575e+00, 8.68645551e+00,
##        8.96658527e+00, 9.24671504e+00, 9.52684480e+00, 9.80697457e+00,
##        1.00871043e+01, 1.03672341e+01, 1.06473639e+01, 1.09274936e+01,
##        1.12076234e+01, 1.14877532e+01, 1.17678829e+01, 1.20480127e+01,
##        1.23281424e+01, 1.26082722e+01, 1.28884020e+01, 1.31685317e+01,
##        1.34486615e+01, 1.37287913e+01, 1.40089210e+01, 1.42890508e+01,
##        1.45691806e+01, 1.48493103e+01, 1.51294401e+01, 1.54095698e+01,
##        1.56896996e+01, 1.59698294e+01, 1.62499591e+01, 1.65300889e+01,
##        1.68102187e+01, 1.70903484e+01, 1.73704782e+01, 1.76506080e+01,
##        1.79307377e+01, 1.82108675e+01, 1.84909973e+01, 1.87711270e+01,
##        1.90512568e+01, 1.93313865e+01, 1.96115163e+01, 1.98916461e+01,
##        2.01717758e+01, 2.04519056e+01, 2.07320354e+01, 2.10121651e+01,
##        2.12922949e+01, 2.15724247e+01, 2.18525544e+01, 2.21326842e+01,
##        2.24128139e+01, 2.26929437e+01, 2.29730735e+01, 2.32532032e+01,
##        2.35333330e+01, 2.38134628e+01, 2.40935925e+01, 2.43737223e+01,
##        2.46538521e+01, 2.49339818e+01, 2.52141116e+01, 2.54942414e+01,
##        2.57743711e+01, 2.60545009e+01, 2.63346306e+01, 2.66147604e+01,
##        2.68948902e+01, 2.71750199e+01, 2.74551497e+01, 2.77352795e+01,
##        2.80154092e+01]), <BarContainer object of 100 artists>)
```

```
axs[0,1].set_title("ALE Inward")
axs[1,1].hist(focus[focus['stimdir']=="outward"]['timefromonset'].values, 100, density=True, facecolor='#593d9c', alpha=0.5)
```

```
## (array([0.05014935, 0.03905863, 0.04291627, 0.04918494, 0.0453273 ,
##        0.05256038, 0.06075787, 0.06316889, 0.05834684, 0.04484509,
##        0.03954083, 0.03423658, 0.02893232, 0.02941452, 0.03278996,
##        0.03230776, 0.02459247, 0.03520099, 0.0270035 , 0.03375437,
##        0.03182555, 0.02989673, 0.03423658, 0.0270035 , 0.02893232,
##        0.03182555, 0.03809422, 0.0366476 , 0.03761201, 0.03809422,
##        0.03520099, 0.03471878, 0.03712981, 0.03423658, 0.03809422,
##        0.03809422, 0.03134334, 0.03761201, 0.03761201, 0.0361654 ,
##        0.0366476 , 0.0361654 , 0.03857642, 0.03954083, 0.03471878,
##        0.03568319, 0.03471878, 0.03954083, 0.04677391, 0.04002304,
##        0.04002304, 0.04291627, 0.04339848, 0.04339848, 0.03905863,
##        0.04966715, 0.03761201, 0.04098745, 0.04050525, 0.05063156,
##        0.03230776, 0.04725612, 0.04725612, 0.03809422, 0.04291627,
##        0.04195186, 0.04243407, 0.04146966, 0.04918494, 0.04291627,
##        0.04098745, 0.03712981, 0.04822053, 0.04002304, 0.05352479,
##        0.0458095 , 0.04050525, 0.04050525, 0.04725612, 0.0544892 ,
##        0.04436289, 0.05063156, 0.04098745, 0.0549714 , 0.04870274,
##        0.04243407, 0.0458095 , 0.04050525, 0.0453273 , 0.03954083,
##        0.04243407, 0.05014935, 0.04822053, 0.05159597, 0.04436289,
##        0.04725612, 0.04388068, 0.0458095 , 0.01928821, 0.00048221]), array([1.05619431e-04, 2.47016618e-01, 4.93927617e-01, 7.40838616e-01,
##        9.87749615e-01, 1.23466061e+00, 1.48157161e+00, 1.72848261e+00,
##        1.97539361e+00, 2.22230461e+00, 2.46921561e+00, 2.71612661e+00,
##        2.96303761e+00, 3.20994860e+00, 3.45685960e+00, 3.70377060e+00,
##        3.95068160e+00, 4.19759260e+00, 4.44450360e+00, 4.69141460e+00,
##        4.93832560e+00, 5.18523659e+00, 5.43214759e+00, 5.67905859e+00,
##        5.92596959e+00, 6.17288059e+00, 6.41979159e+00, 6.66670259e+00,
##        6.91361359e+00, 7.16052459e+00, 7.40743558e+00, 7.65434658e+00,
##        7.90125758e+00, 8.14816858e+00, 8.39507958e+00, 8.64199058e+00,
##        8.88890158e+00, 9.13581258e+00, 9.38272357e+00, 9.62963457e+00,
##        9.87654557e+00, 1.01234566e+01, 1.03703676e+01, 1.06172786e+01,
##        1.08641896e+01, 1.11111006e+01, 1.13580116e+01, 1.16049226e+01,
##        1.18518336e+01, 1.20987446e+01, 1.23456556e+01, 1.25925666e+01,
##        1.28394776e+01, 1.30863886e+01, 1.33332996e+01, 1.35802106e+01,
##        1.38271216e+01, 1.40740326e+01, 1.43209436e+01, 1.45678545e+01,
##        1.48147655e+01, 1.50616765e+01, 1.53085875e+01, 1.55554985e+01,
##        1.58024095e+01, 1.60493205e+01, 1.62962315e+01, 1.65431425e+01,
##        1.67900535e+01, 1.70369645e+01, 1.72838755e+01, 1.75307865e+01,
##        1.77776975e+01, 1.80246085e+01, 1.82715195e+01, 1.85184305e+01,
##        1.87653415e+01, 1.90122525e+01, 1.92591635e+01, 1.95060745e+01,
##        1.97529855e+01, 1.99998965e+01, 2.02468075e+01, 2.04937185e+01,
##        2.07406295e+01, 2.09875405e+01, 2.12344515e+01, 2.14813625e+01,
##        2.17282735e+01, 2.19751845e+01, 2.22220955e+01, 2.24690065e+01,
##        2.27159175e+01, 2.29628285e+01, 2.32097395e+01, 2.34566505e+01,
##        2.37035615e+01, 2.39504725e+01, 2.41973835e+01, 2.44442945e+01,
##        2.46912055e+01]), <BarContainer object of 100 artists>)
```

```
axs[1,1].set_title("ALE Outward")

focus= exp2[exp2['eyes']=="PLE"]
axs[0,2].hist(focus[focus['stimdir']=="inward"]['timefromonset'].values, 100, density=True, facecolor='#593d9c', alpha=0.5)
```

```
## (array([0.04655448, 0.03542189, 0.04048216, 0.04655448, 0.04554243,
##        0.05515694, 0.05313283, 0.04908462, 0.04453038, 0.03643394,
##        0.03896408, 0.04200024, 0.03845805, 0.03997613, 0.04453038,
##        0.0318797 , 0.03795202, 0.03795202, 0.03542189, 0.04048216,
##        0.04250627, 0.04959065, 0.03845805, 0.04200024, 0.04402435,
##        0.05161475, 0.06578351, 0.05971118, 0.06730159, 0.05566297,
##        0.05262681, 0.04351832, 0.03693997, 0.03036162, 0.02934957,
##        0.02985559, 0.02884354, 0.02530135, 0.02580738, 0.02783148,
##        0.0242893 , 0.02530135, 0.02530135, 0.02884354, 0.0318797 ,
##        0.03289175, 0.0263134 , 0.03289175, 0.02732546, 0.03339778,
##        0.02530135, 0.03896408, 0.03491586, 0.037446  , 0.03643394,
##        0.03947011, 0.03086765, 0.037446  , 0.03440984, 0.03440984,
##        0.04098819, 0.04149421, 0.04149421, 0.04149421, 0.04857859,
##        0.04959065, 0.03592792, 0.04301229, 0.03896408, 0.04402435,
##        0.04554243, 0.04250627, 0.03947011, 0.04655448, 0.04604846,
##        0.04807256, 0.04453038, 0.04807256, 0.03845805, 0.05009667,
##        0.04756654, 0.04604846, 0.04604846, 0.0450364 , 0.0506027 ,
##        0.04756654, 0.04706051, 0.04453038, 0.04706051, 0.03845805,
##        0.04706051, 0.04807256, 0.04604846, 0.03896408, 0.04402435,
##        0.04756654, 0.05009667, 0.04756654, 0.01922903, 0.00101205]), array([5.06877899e-03, 2.51967726e-01, 4.98866673e-01, 7.45765619e-01,
##        9.92664566e-01, 1.23956351e+00, 1.48646246e+00, 1.73336141e+00,
##        1.98026035e+00, 2.22715930e+00, 2.47405825e+00, 2.72095719e+00,
##        2.96785614e+00, 3.21475509e+00, 3.46165403e+00, 3.70855298e+00,
##        3.95545193e+00, 4.20235087e+00, 4.44924982e+00, 4.69614877e+00,
##        4.94304771e+00, 5.18994666e+00, 5.43684561e+00, 5.68374455e+00,
##        5.93064350e+00, 6.17754245e+00, 6.42444139e+00, 6.67134034e+00,
##        6.91823929e+00, 7.16513824e+00, 7.41203718e+00, 7.65893613e+00,
##        7.90583508e+00, 8.15273402e+00, 8.39963297e+00, 8.64653192e+00,
##        8.89343086e+00, 9.14032981e+00, 9.38722876e+00, 9.63412770e+00,
##        9.88102665e+00, 1.01279256e+01, 1.03748245e+01, 1.06217235e+01,
##        1.08686224e+01, 1.11155214e+01, 1.13624203e+01, 1.16093193e+01,
##        1.18562182e+01, 1.21031172e+01, 1.23500161e+01, 1.25969151e+01,
##        1.28438140e+01, 1.30907130e+01, 1.33376119e+01, 1.35845109e+01,
##        1.38314098e+01, 1.40783087e+01, 1.43252077e+01, 1.45721066e+01,
##        1.48190056e+01, 1.50659045e+01, 1.53128035e+01, 1.55597024e+01,
##        1.58066014e+01, 1.60535003e+01, 1.63003993e+01, 1.65472982e+01,
##        1.67941972e+01, 1.70410961e+01, 1.72879951e+01, 1.75348940e+01,
##        1.77817929e+01, 1.80286919e+01, 1.82755908e+01, 1.85224898e+01,
##        1.87693887e+01, 1.90162877e+01, 1.92631866e+01, 1.95100856e+01,
##        1.97569845e+01, 2.00038835e+01, 2.02507824e+01, 2.04976814e+01,
##        2.07445803e+01, 2.09914793e+01, 2.12383782e+01, 2.14852771e+01,
##        2.17321761e+01, 2.19790750e+01, 2.22259740e+01, 2.24728729e+01,
##        2.27197719e+01, 2.29666708e+01, 2.32135698e+01, 2.34604687e+01,
##        2.37073677e+01, 2.39542666e+01, 2.42011656e+01, 2.44480645e+01,
##        2.46949635e+01]), <BarContainer object of 100 artists>)
```

```
axs[0,2].set_title("PLE Inward")
axs[1,2].hist(focus[focus['stimdir']=="outward"]['timefromonset'].values, 100, density=True, facecolor='#593d9c', alpha=0.5)
```

```
## (array([0.04671415, 0.04767733, 0.04719574, 0.05249323, 0.05249323,
##        0.05008528, 0.06501455, 0.05634594, 0.06308819, 0.06405137,
##        0.06983044, 0.06068024, 0.06356978, 0.05586435, 0.04526939,
##        0.03997191, 0.03611919, 0.04141667, 0.04767733, 0.04430621,
##        0.04575098, 0.04141667, 0.05201164, 0.0669409 , 0.07031203,
##        0.05730912, 0.04719574, 0.03852714, 0.03419283, 0.02263469,
##        0.0221531 , 0.0221531 , 0.01589244, 0.0221531 , 0.02311628,
##        0.01974516, 0.0221531 , 0.02070834, 0.02793217, 0.02263469,
##        0.02552423, 0.02407946, 0.02600582, 0.026969  , 0.02407946,
##        0.02841376, 0.02937694, 0.03515601, 0.0313033 , 0.02985853,
##        0.03515601, 0.03178489, 0.03322966, 0.03804555, 0.03371125,
##        0.03660078, 0.0356376 , 0.03756396, 0.03708237, 0.04334303,
##        0.04526939, 0.0356376 , 0.03804555, 0.04286144, 0.03804555,
##        0.04189826, 0.03419283, 0.03660078, 0.03900873, 0.04334303,
##        0.03804555, 0.04093508, 0.04960369, 0.04237985, 0.04526939,
##        0.04382462, 0.05249323, 0.03708237, 0.05393799, 0.05056687,
##        0.05056687, 0.05104846, 0.0447878 , 0.04575098, 0.04286144,
##        0.04189826, 0.04623257, 0.04864051, 0.04575098, 0.04334303,
##        0.01107655, 0.00048159, 0.00048159, 0.        , 0.00096318,
##        0.        , 0.        , 0.00096318, 0.        , 0.00048159]), array([6.82592392e-04, 2.70072632e-01, 5.39462671e-01, 8.08852711e-01,
##        1.07824275e+00, 1.34763279e+00, 1.61702283e+00, 1.88641287e+00,
##        2.15580291e+00, 2.42519295e+00, 2.69458299e+00, 2.96397303e+00,
##        3.23336307e+00, 3.50275311e+00, 3.77214314e+00, 4.04153318e+00,
##        4.31092322e+00, 4.58031326e+00, 4.84970330e+00, 5.11909334e+00,
##        5.38848338e+00, 5.65787342e+00, 5.92726346e+00, 6.19665350e+00,
##        6.46604354e+00, 6.73543358e+00, 7.00482362e+00, 7.27421366e+00,
##        7.54360370e+00, 7.81299374e+00, 8.08238378e+00, 8.35177382e+00,
##        8.62116385e+00, 8.89055389e+00, 9.15994393e+00, 9.42933397e+00,
##        9.69872401e+00, 9.96811405e+00, 1.02375041e+01, 1.05068941e+01,
##        1.07762842e+01, 1.10456742e+01, 1.13150642e+01, 1.15844543e+01,
##        1.18538443e+01, 1.21232344e+01, 1.23926244e+01, 1.26620144e+01,
##        1.29314045e+01, 1.32007945e+01, 1.34701846e+01, 1.37395746e+01,
##        1.40089646e+01, 1.42783547e+01, 1.45477447e+01, 1.48171348e+01,
##        1.50865248e+01, 1.53559148e+01, 1.56253049e+01, 1.58946949e+01,
##        1.61640850e+01, 1.64334750e+01, 1.67028650e+01, 1.69722551e+01,
##        1.72416451e+01, 1.75110352e+01, 1.77804252e+01, 1.80498152e+01,
##        1.83192053e+01, 1.85885953e+01, 1.88579854e+01, 1.91273754e+01,
##        1.93967654e+01, 1.96661555e+01, 1.99355455e+01, 2.02049356e+01,
##        2.04743256e+01, 2.07437156e+01, 2.10131057e+01, 2.12824957e+01,
##        2.15518857e+01, 2.18212758e+01, 2.20906658e+01, 2.23600559e+01,
##        2.26294459e+01, 2.28988359e+01, 2.31682260e+01, 2.34376160e+01,
##        2.37070061e+01, 2.39763961e+01, 2.42457861e+01, 2.45151762e+01,
##        2.47845662e+01, 2.50539563e+01, 2.53233463e+01, 2.55927363e+01,
##        2.58621264e+01, 2.61315164e+01, 2.64009065e+01, 2.66702965e+01,
##        2.69396865e+01]), <BarContainer object of 100 artists>)
```

```
axs[1,2].set_title("PLE Outward")
```

We can only see an effect on the outward moving dots, however when
looking at the graph we can appreciate an end peak towards high angles
for the inward moving stimuli. This must be an effect of appearance, as
stated initially. For inward moving stimuli the appearance is at around
90 deg, where actually only some eyes can see (PLE). Outward stimuli
appear at 0 degrees, but for PLE appear at 50deg, where they disappear
for ALE. Overall outward stimuli follow a more gradual appearance for
our experimental condition.

Overall, this is telling me that I need to select experiments’
section where to look for behaviour. Stimuli should be in their
appearing phase, for each condition. this poses our selection this way:
- ALE: inward moving stimuli, at around 60-40 degrees - PLE: outward
moving stimuli, at around the same, slightly moved due to the shifted
start of the field, 70-50 - Control: more complicated. Should be at
around 90 and around 0. however at around 90 we will only observe
stimuli seen by PLE exclusively. at 0, ALE exclusively and moreover
pivots will not happen. we could select 70-40 to get all of the selected
field of the two above, but we risk getting the effect muddied by the
fact that stimuli appeared long before. I will keep all the data, but I
don’t expect to observe big effects.

Now, to cutting the table:

```
ALE <- subset(exp2, exp2$eyes == 'ALE')
PLE <- subset(exp2, exp2$eyes == 'PLE')
Control <- subset(exp2, exp2$eyes == 'Control')

ALE <- subset(ALE, ALE$stimdir == 'inward')
ALE <- subset(ALE, ALE$stimpos < 60)
ALE <- subset(ALE, ALE$stimpos > 40)

PLE <- subset(PLE, PLE$stimdir == 'outward')
PLE <- subset(PLE, PLE$stimpos < 70)
PLE <- subset(PLE, PLE$stimpos > 50)


exp2cut <- rbind(ALE, PLE, Control)
exp2cut <- subset(exp2cut, exp2cut$stimpres ==1)
```

##### Main Analysis

Now to see the preference.

```
hist(exp2$dirval_deg_s, breaks=100)
```

```
mmain <- glmmTMB(dirval_deg_s~eyes*cond + (cond|subj),
                 data=exp2cut, family= gaussian(),
                 control=glmmTMBControl(optCtrl = list(iter.max = 300000, eval.max = 400000)))
```

```
## Warning in fitTMB(TMBStruc): Model convergence problem; non-positive-definite
## Hessian matrix. See vignette('troubleshooting')
```

```
diagnose(mmain)
```

```
## Unusually large Z-statistics (|x|>5):
## 
## d~(Intercept) 
##      574.4223 
## 
## Large Z-statistics (estimate/std err) suggest a *possible* failure of
## the Wald approximation - often also associated with parameters that are
## at or near the edge of their range (e.g. random-effects standard
## deviations approaching 0).  (Alternately, they may simply represent
## very well-estimated parameters; intercepts of non-centered models may
## fall in this category.) While the Wald p-values and standard errors
## listed in summary() may be unreliable, profile confidence intervals
## (see ?confint.glmmTMB) and likelihood ratio test p-values derived by
## comparing models (e.g. ?drop1) are probably still OK.  (Note that the
## LRT is conservative when the null value is on the boundary, e.g. a
## variance or zero-inflation value of 0 (Self and Liang 1987; Stram and
## Lee 1994; Goldman and Whelan 2000); in simple cases these p-values are
## approximately twice as large as they should be.)
## 
## 
## Non-positive definite (NPD) Hessian
## 
## The Hessian matrix represents the curvature of the log-likelihood
## surface at the maximum likelihood estimate (MLE) of the parameters (its
## inverse is the estimate of the parameter covariance matrix).  A
## non-positive-definite Hessian means that the likelihood surface is
## approximately flat (or upward-curving) at the MLE, which means the
## model is overfitted or poorly posed in some way. NPD Hessians are often
## associated with extreme parameter estimates.
## 
## 
## parameters with non-finite standard deviations:
## theta_cond|subj.2
## 
## 
## 
## recomputing Hessian via Richardson extrapolation. If this is too slow, consider setting check_hessian = FALSE 
## 
## The next set of diagnostics attempts to determine which elements of the
## Hessian are causing the non-positive-definiteness.  Components with
## very small eigenvalues represent 'flat' directions, i.e., combinations
## of parameters for which the data may contain very little information.
## So-called 'bad elements' represent the dominant components (absolute
## values >0.01) of the eigenvectors corresponding to the 'flat'
## directions
## 
## 
## maximum Hessian eigenvalue = 4.45e+03 
## Hessian eigenvalue 7 = 0.0167 (relative val = 3.75e-06) 
##    bad elements: (Intercept) eyesControl eyesPLE condsilh-ellipse eyesControl:condsilh-ellipse eyesPLE:condsilh-ellipse 
## Hessian eigenvalue 8 = 0.000122 (relative val = 2.74e-08) 
##    bad elements: theta_cond|subj.1 theta_cond|subj.2 theta_cond|subj.3 
## Hessian eigenvalue 9 = 6.41e-05 (relative val = 1.44e-08) 
##    bad elements: theta_cond|subj.1 theta_cond|subj.2 theta_cond|subj.3 
## Hessian eigenvalue 10 = -3e-05 (relative val = -6.75e-09) 
##    bad elements: theta_cond|subj.1 theta_cond|subj.2 theta_cond|subj.3
```

the model is overfitted. I will drop cond from the random effect as
the variance is apparently explained by the fixed version of the effect
itself.

```
mmain <- glmmTMB(dirval_deg_s~eyes*cond + (1|subj),
                 data=exp2cut, family= gaussian(),
                 control=glmmTMBControl(optCtrl = list(iter.max = 300000, eval.max = 400000)))

simres <- simulateResiduals(mmain)
plot(simres)
```

Even though Dharma gives error, it is clearly a gaussian from the
histogram

```
Anova(mmain)
```

```
## Warning in printHypothesis(L, rhs, names(b)): one or more coefficients in the hypothesis include
##      arithmetic operators in their names;
##   the printed representation of the hypothesis will be omitted

## Warning in printHypothesis(L, rhs, names(b)): one or more coefficients in the hypothesis include
##      arithmetic operators in their names;
##   the printed representation of the hypothesis will be omitted

## Warning in printHypothesis(L, rhs, names(b)): one or more coefficients in the hypothesis include
##      arithmetic operators in their names;
##   the printed representation of the hypothesis will be omitted
```

```
## Analysis of Deviance Table (Type II Wald chisquare tests)
## 
## Response: dirval_deg_s
##            Chisq Df Pr(>Chisq)  
## eyes      3.6401  2    0.16202  
## cond      6.2842  1    0.01218 *
## eyes:cond 3.4059  2    0.18214  
## ---
## Signif. codes:  0 '***' 0.001 '**' 0.01 '*' 0.05 '.' 0.1 ' ' 1
```

```
e <- emmeans(mmain, ~cond*eyes, type='response')
e
```

```
##  cond         eyes     emmean   SE   df lower.CL upper.CL
##  bio-rand     ALE      7.5723 2.52 8884     2.63   12.515
##  silh-ellipse ALE     -1.9663 2.32 8884    -6.51    2.575
##  bio-rand     Control  0.2180 1.44 8884    -2.61    3.045
##  silh-ellipse Control -2.0592 1.37 8884    -4.74    0.621
##  bio-rand     PLE      3.1869 3.17 8884    -3.04    9.410
##  silh-ellipse PLE     -0.0123 2.55 8884    -5.00    4.977
## 
## Confidence level used: 0.95
```

```
test(e, adjust='bonferroni')
```

```
##  cond         eyes     emmean   SE   df t.ratio p.value
##  bio-rand     ALE      7.5723 2.52 8884   3.003  0.0161
##  silh-ellipse ALE     -1.9663 2.32 8884  -0.849  1.0000
##  bio-rand     Control  0.2180 1.44 8884   0.151  1.0000
##  silh-ellipse Control -2.0592 1.37 8884  -1.506  0.7926
##  bio-rand     PLE      3.1869 3.17 8884   1.004  1.0000
##  silh-ellipse PLE     -0.0123 2.55 8884  -0.005  1.0000
## 
## P value adjustment: bonferroni method for 6 tests
```

```
write.csv(as.data.frame(e), paste0(path, "binomialresults.csv"))
```

```
toplot = pd.read_csv(path+'binomialresults.csv')


ALE = toplot[toplot['eyes']=='ALE']
PLE = toplot[toplot['eyes']=='PLE']
Control = toplot[toplot['eyes']=='Control']

fig, axs = plt.subplots(3, 1)

for n, tab in enumerate([Control, ALE, PLE]):
        axs[n].axvline(x=tab[tab['cond']=='bio-rand'].emmean.values[0], c='#63146e', linewidth=5)
        axs[n].axvspan(tab[tab['cond']=='bio-rand'].emmean.values[0]-tab.SE.values[0],
                          tab[tab['cond']=='bio-rand'].emmean.values[0]+tab.SE.values[0], color='#63146e', alpha=0.8)

        axs[n].axvline(x=tab[tab['cond']=='silh-ellipse'].emmean.values[0], c='#806016', linewidth=5)
        axs[n].axvspan(tab[tab['cond']=='silh-ellipse'].emmean.values[0]-tab.SE.values[0],
                          tab[tab['cond']=='silh-ellipse'].emmean.values[0]+tab.SE.values[0], color='#daa520', alpha=0.8)

        axs[n].xaxis.set_tick_params(labelsize=14)
        axs[n].yaxis.set_tick_params(labelsize=14)
        axs[n].set_xticks(np.arange(-12, 14, 2))
        axs[n].axvline(x=0, c='black', linestyle='dotted')
        axs[n].set_ylabel(tab.eyes.values[0], fontsize=10)
```

For the control condition the choice is random. this is probably due
to what has been said earlier: the stimuli appear at different sections,
where only one eye pair can see them. We cannot draw conclusions for
PLE, as they show no significant result either. Crucially, ALE show a
preference for the biological, for point light displays but not for
silh

##### Is there a side bias?

We need to test whether there is a pre-existing side bias. The
position of the stimuli is counterbalanced across trials, so even if
there is a side bias it will not influence the results of the
experiment, but still it will be useful to know.

```
mside <- glmmTMB(dirval_deg_s_lr~eyes*cond + (1|subj/trialn),
                 data = exp2cut, family = gaussian,
                control=glmmTMBControl(optCtrl = list(iter.max = 30000, eval.max = 40000)))

Anova(mside)
```

```
## Warning in printHypothesis(L, rhs, names(b)): one or more coefficients in the hypothesis include
##      arithmetic operators in their names;
##   the printed representation of the hypothesis will be omitted

## Warning in printHypothesis(L, rhs, names(b)): one or more coefficients in the hypothesis include
##      arithmetic operators in their names;
##   the printed representation of the hypothesis will be omitted

## Warning in printHypothesis(L, rhs, names(b)): one or more coefficients in the hypothesis include
##      arithmetic operators in their names;
##   the printed representation of the hypothesis will be omitted
```

```
## Analysis of Deviance Table (Type II Wald chisquare tests)
## 
## Response: dirval_deg_s_lr
##            Chisq Df Pr(>Chisq)   
## eyes      4.4758  2   0.106680   
## cond      7.5066  1   0.006147 **
## eyes:cond 7.8143  2   0.020098 * 
## ---
## Signif. codes:  0 '***' 0.001 '**' 0.01 '*' 0.05 '.' 0.1 ' ' 1
```

```
e<-emmeans(mside, ~cond, type='response')
```

```
## NOTE: Results may be misleading due to involvement in interactions
```

```
test(e, adjust='bonferroni')
```

```
##  cond         emmean   SE   df t.ratio p.value
##  bio-rand      -3.28 1.77 8883  -1.856  0.1271
##  silh-ellipse   4.43 1.57 8883   2.823  0.0095
## 
## Results are averaged over the levels of: eyes 
## P value adjustment: bonferroni method for 2 tests
```

There is an effect of condition on the side bias. Indeed, there seems
to be a mild right preference but only for the shil-ellipse condition.
As the stimuli are balanced, this is not a problem for the result, but
for a possible effect weakening.

##### Are there differences between the sexes?

It may be interesting to see whether there are differences in the
performance between male, females and juveniles (unsexed) spiders. We
decided not to include it in the main analysis as the subjects were not
paired by sex:

```
summary(exp2$sex[!duplicated(exp2$subj)])
```

```
##  f  j  m 
## 89 75 15
```

As we mostly have female and juvenile spiders, a lack of a
significant preference in males should be taken with caution, as it may
just be due to a low N. Moreover, adding sex as a predictor will divide
the sample and force us to perform many multiple comparisons, increasing
the effect of a bonferroni correction and decreasing our statistical
power.

Keeping in mind these premises, we will redo the main analysis
including sex as a predictor and discuss the results, without giving too
much weight to the p-values themselves: even if we are unable to provide
a definitive result about it, it may inform other scientists about
further experiments.

We will skip the checks about the data distribution, since our
dependent variable remained the same.

```
msex <- glmmTMB(dirval_deg_s~cond*eyes*sex + (1|subj/trialn),
                 data=exp2cut, family= gaussian(),
                 control=glmmTMBControl(optCtrl = list(iter.max = 30000, eval.max = 40000)))

Anova(msex)
```

```
## Warning in printHypothesis(L, rhs, names(b)): one or more coefficients in the hypothesis include
##      arithmetic operators in their names;
##   the printed representation of the hypothesis will be omitted

## Warning in printHypothesis(L, rhs, names(b)): one or more coefficients in the hypothesis include
##      arithmetic operators in their names;
##   the printed representation of the hypothesis will be omitted

## Warning in printHypothesis(L, rhs, names(b)): one or more coefficients in the hypothesis include
##      arithmetic operators in their names;
##   the printed representation of the hypothesis will be omitted

## Warning in printHypothesis(L, rhs, names(b)): one or more coefficients in the hypothesis include
##      arithmetic operators in their names;
##   the printed representation of the hypothesis will be omitted

## Warning in printHypothesis(L, rhs, names(b)): one or more coefficients in the hypothesis include
##      arithmetic operators in their names;
##   the printed representation of the hypothesis will be omitted

## Warning in printHypothesis(L, rhs, names(b)): one or more coefficients in the hypothesis include
##      arithmetic operators in their names;
##   the printed representation of the hypothesis will be omitted

## Warning in printHypothesis(L, rhs, names(b)): one or more coefficients in the hypothesis include
##      arithmetic operators in their names;
##   the printed representation of the hypothesis will be omitted
```

```
## Analysis of Deviance Table (Type II Wald chisquare tests)
## 
## Response: dirval_deg_s
##                Chisq Df Pr(>Chisq)  
## cond          5.8171  1    0.01587 *
## eyes          2.9690  2    0.22661  
## sex           1.3333  2    0.51343  
## cond:eyes     3.5647  2    0.16824  
## cond:sex      1.7768  2    0.41131  
## eyes:sex      1.9132  4    0.75172  
## cond:eyes:sex 3.0849  4    0.54371  
## ---
## Signif. codes:  0 '***' 0.001 '**' 0.01 '*' 0.05 '.' 0.1 ' ' 1
```

As shown here, even when adding sex as a predictor, our main effects
of condition remain.

Regarding sex, there seems to be no main effect, meaning that
females, males and juveniles all respond with overall similar frequency.
Moreover, no two-ways interaction with sex result significant.

We will proceed anyway with a post hoc analysis

```
e <- emmeans(msex, ~cond*eyes*sex, type='response')


test(e, adjust='bonferroni', simple=sex)
```

```
##  cond         eyes    sex  emmean    SE   df t.ratio p.value
##  bio-rand     ALE     f     5.381  3.36 8871   1.600  1.0000
##  silh-ellipse ALE     f    -2.356  3.40 8871  -0.693  1.0000
##  bio-rand     Control f    -0.188  1.72 8871  -0.109  1.0000
##  silh-ellipse Control f    -0.948  1.75 8871  -0.543  1.0000
##  bio-rand     PLE     f     1.170  4.37 8871   0.268  1.0000
##  silh-ellipse PLE     f    -1.444  3.51 8871  -0.412  1.0000
##  bio-rand     ALE     j    10.986  4.07 8871   2.701  0.1246
##  silh-ellipse ALE     j    -2.109  3.46 8871  -0.610  1.0000
##  bio-rand     Control j     0.331  2.92 8871   0.113  1.0000
##  silh-ellipse Control j    -0.919  2.60 8871  -0.353  1.0000
##  bio-rand     PLE     j     6.328  5.43 8871   1.166  1.0000
##  silh-ellipse PLE     j     0.772  4.02 8871   0.192  1.0000
##  bio-rand     ALE     m     6.097 10.83 8871   0.563  1.0000
##  silh-ellipse ALE     m     0.831  7.82 8871   0.106  1.0000
##  bio-rand     Control m     4.888  6.14 8871   0.796  1.0000
##  silh-ellipse Control m   -10.980  4.09 8871  -2.683  0.1314
##  bio-rand     PLE     m     3.123  8.81 8871   0.355  1.0000
##  silh-ellipse PLE     m     6.024  9.43 8871   0.639  1.0000
## 
## P value adjustment: bonferroni method for 18 tests
```

This is not worth discussing, as there are too many comparisons for
it to be meaningful. Is kept for completeness.

### Session info

```
## R version 4.3.1 (2023-06-16)
## Platform: x86_64-pc-linux-gnu (64-bit)
## Running under: EndeavourOS
## 
## Matrix products: default
## BLAS:   /usr/lib/libblas.so.3.11.0 
## LAPACK: /home/massimodeagro/anaconda3/envs/DataAnalysis/lib/libmkl_rt.so.2;  LAPACK version 3.10.1
## 
## locale:
##  [1] LC_CTYPE=it_IT.UTF-8       LC_NUMERIC=C              
##  [3] LC_TIME=it_IT.UTF-8        LC_COLLATE=it_IT.UTF-8    
##  [5] LC_MONETARY=it_IT.UTF-8    LC_MESSAGES=it_IT.UTF-8   
##  [7] LC_PAPER=it_IT.UTF-8       LC_NAME=C                 
##  [9] LC_ADDRESS=C               LC_TELEPHONE=C            
## [11] LC_MEASUREMENT=it_IT.UTF-8 LC_IDENTIFICATION=C       
## 
## time zone: Europe/Rome
## tzcode source: system (glibc)
## 
## attached base packages:
## [1] stats     graphics  grDevices utils     datasets  methods   base     
## 
## other attached packages:
## [1] reticulate_1.28 ggplot2_3.4.2   emmeans_1.8.6   DHARMa_0.4.6   
## [5] car_3.1-2       carData_3.0-5   glmmTMB_1.1.7   readODS_2.0.0  
## 
## loaded via a namespace (and not attached):
##  [1] gtable_0.3.3        TMB_1.9.4           xfun_0.39          
##  [4] bslib_0.4.2         lattice_0.21-8      numDeriv_2016.8-1.1
##  [7] vctrs_0.6.2         tools_4.3.1         generics_0.1.3     
## [10] parallel_4.3.1      tibble_3.2.1        fansi_1.0.4        
## [13] highr_0.10          pkgconfig_2.0.3     KernSmooth_2.23-21 
## [16] Matrix_1.5-4.1      lifecycle_1.0.3     compiler_4.3.1     
## [19] munsell_0.5.0       gap.datasets_0.0.5  codetools_0.2-19   
## [22] httpuv_1.6.11       htmltools_0.5.5     sass_0.4.6         
## [25] yaml_2.3.7          later_1.3.1         pillar_1.9.0       
## [28] nloptr_2.0.3        jquerylib_0.1.4     ellipsis_0.3.2     
## [31] MASS_7.3-60         cachem_1.0.8        iterators_1.0.14   
## [34] boot_1.3-28.1       abind_1.4-5         foreach_1.5.2      
## [37] mime_0.12           nlme_3.1-162        tidyselect_1.2.0   
## [40] digest_0.6.31       mvtnorm_1.1-3       dplyr_1.1.2        
## [43] splines_4.3.1       rprojroot_2.0.3     fastmap_1.1.1      
## [46] grid_4.3.1          here_1.0.1          colorspace_2.1-0   
## [49] cli_3.6.1           magrittr_2.0.3      utf8_1.2.3         
## [52] withr_2.5.0         promises_1.2.0.1    scales_1.2.1       
## [55] estimability_1.4.1  rmarkdown_2.21      lme4_1.1-33        
## [58] png_0.1-8           shiny_1.7.4         evaluate_0.21      
## [61] knitr_1.43          gap_1.5-1           doParallel_1.0.17  
## [64] mgcv_1.8-42         rlang_1.1.1         Rcpp_1.0.10        
## [67] xtable_1.8-4        glue_1.6.2          qgam_1.3.4         
## [70] minqa_1.2.5         jsonlite_1.8.4      R6_2.5.1           
## [73] plyr_1.8.8
```
